# ciRS-7 and miR-7 regulate ischemia induced neuronal death via glutamatergic signaling

**DOI:** 10.1101/2023.01.24.525136

**Authors:** Flavia Scoyni, Valeriia Sitnikova, Luca Giudice, Paula Korhonen, Davide M Trevisan, Ana Hernandez de Sande, Mireia Gomez-Budia, Raisa Giniatullina, Irene F Ugidos, Hiramani Dhungana, Cristiana Pistono, Nea Korvenlaita, Nelli-Noora Välimäki, Salla M Kangas, Anniina E Hiltunen, Emma Gribchenko, Minna U Kaikkonen-Määttä, Jari Koistinaho, Seppo Ylä-Herttuala, Reetta Hinttala, Morten T Venø, Junyi Su, Markus Stoffel, Anne Schaefer, Nikolaus Rajewsky, Jørgen Kjems, Mary P LaPierre, Monika Piwecka, Jukka Jolkkonen, Rashid Giniatullin, Thomas B Hansen, Tarja Malm

## Abstract

Brain functionality relies on finely tuned regulation of gene expression by networks of non-coding RNAs (ncRNAs) such as the one composed by the circular RNA ciRS-7 (also known as CDR1as), the microRNA miR-7 and the long non-coding RNA Cyrano. Here we describe ischemia induced alterations in the ncRNA network both *in vitro* and *in vivo* and in transgenic mice lacking ciRS-7 or miR-7. Our data show that cortical neurons downregulate ciRS-7 and Cyrano and upregulate miR-7 expression upon ischemic insults. Mice lacking ciRS-7 show reduced lesion size and motor impairment, whilst the absence of miR-7 alone leads to an increase in the ischemia induced neuronal death. Moreover, miR-7 levels in pyramidal excitatory neurons regulate dendrite morphology and glutamatergic signaling suggesting a potential molecular link to the *in vivo* phenotype. Our data reveal that ciRS-7 and miR-7 contribute to the outcome of ischemic stroke and shed new light into the pathophysiological roles of intracellular networks of non-coding RNAs in the brain.

## INTRODUCTION

The intricate functionality of the brain relies on precisely regulated gene expression, also mediated by non-coding RNAs (ncRNAs), molecules abundant in the brain linked to the increased cognitive complexity in human^1^. MicroRNAs (miRNAs), short (∼22 nucleotides) ncRNAs, post-transcriptionally regulate messenger RNA (mRNA) expression by binding short complementary sequences (seed)^2^ and triggering mRNA decay or inhibition of translation^3^. Long non-coding RNAs (lncRNAs), RNA molecules longer than 200 nucleotides, regulate gene expression by interacting with other ncRNAs, including miRNAs, or proteins^4^. Circular RNAs (circRNAs), a novel class of ncRNAs, modulate gene expression also by interacting with miRNAs, hence affecting miRNAs activity on mRNA targets^5^. CircRNAs result from backsplicing of linear transcripts, an uncanonical splicing event in which a 5’ splice site is spliced with the 3’ splice site of the upstream exon^6^, forming a circular molecule.

These ncRNAs independently control cellular function by regulating the expression of protein-coding genes, but also interact with each other. In the brain-specific ncRNA network involving miR-7, miR-671, the lncRNA Cyrano and the circRNA ciRS-7 (also known as CDR1as)^5,7–9^, ciRS-7 is suggested to stabilize and promote miR-7 targeting^8,9^, while Cyrano triggers miR-7 degradation via target RNA– directed miRNA degradation (TDMD)^8^ with a nearly perfectly complementary binding site. Analysis through gain and loss of function experiments and knock-out animals revealed that Cyrano and ciRS-7 bind miR-7, regulating its expression^5,8,9^ and that Cyrano TDMD effect on miR-7 affects ciRS-7 abundance and localization, indirectly affecting the gene expression of miR-7 targets^5,8–10^. Nonetheless, the physiological purpose of this network in the brain remains unknown, including lack of information on its role in pathophysiological conditions.

Ischemic stroke, induced by occlusion of one of the major cerebral arteries, leads nutrients and oxygen deprivation in the brain parenchyma inducing cell death. Lack of energy and disruption in the ion balance cause an uncontrolled release of glutamate in excitatory neurons, leading to excitotoxicity^11^, oxidative stress, necrosis, and apoptosis^12^. Various miRNAs and lncRNAs have been implicated in regulating oxidative stress response and glutamate excitotoxicity, impacting stroke outcomes^13^. Sustained stress prompts miRNAs to facilitate adaptive switches in gene expression program^14^. The efficiency of the miRNA-mediated response depends on miRNAs availability to interact (expression, localization, activity) and the amount of mRNA targets possessing a Mirna Recognition Element (MRE), creating a miRNA-specific threshold^15^. Sudden changes in MRE-containing sequences can disrupt miRNA targeting and derepress specific targets. This crosstalk led to theorize that in physiological conditions a large number of MRE sequences compete for the binding of the miRNA^16^. Organisms exploit this phenomenon during stress by regulating miRNA activity through transcripts with varying binding strength or MRE abundance following a process called target mimicry^17^.

Recent studies emphasize a strong connection between circRNAs and reactive oxygen species production^18^. Despite individual ncRNAs being studied in the context of ischemic stroke^13^, a comprehensive analysis of the regulatory role of ncRNAs network in stroke is still lacking. In this study, we identified ciRS-7, brain enriched circRNA, ciRS-7, among seven circRNAs deregulated in ischemia-related conditions *in vitro* and *in vivo*. Expanding our analysis to other ncRNAs species, we identified changes in miR-7 and Cyrano, part of the same regulatory circuitry as ciRS-7. Using ciRS-7 and miR-7 knock-out mouse models, we revealed the potential role of ciRS-7 in preventing miR-7-mediated regulation of the target mRNAs, unveiling the contribution of ciRS-7 molecular network in preventing stroke-induced cell death.

## RESULTS

### Oxygen and glucose deprivation induces changes in the expression of circular RNAs

Given the association of circRNAs with oxidative stress response^18^ we evaluated their potential deregulation in conditions mimicking ischemic stroke *in vitro*. We cultured murine cortical neurons isolated from embryonic day fifteen (E15) cortices, subjected them to oxygen and glucose deprivation (OGD) for 12 hours (Supplementary Figure S1A) and performed total RNA sequencing.

A deconvolution analysis on single-cell RNA-seq dataset of embryonic mouse brain (E14.5^19^) confirmed that our culture is representative of the murine cortex, identifying interneurons (Int) and pyramidal neurons (Layer V-VI) as the predominant cell populations (Figure 1A). Functional validation using calcium imaging recording upon GABA and glutamate stimulation revealed that our culture primarily consists of glutamatergic excitatory neurons, as 98% of cells in our culture responded to glutamate and 36% to GABA (Figure 1B).

**Figure 1.**
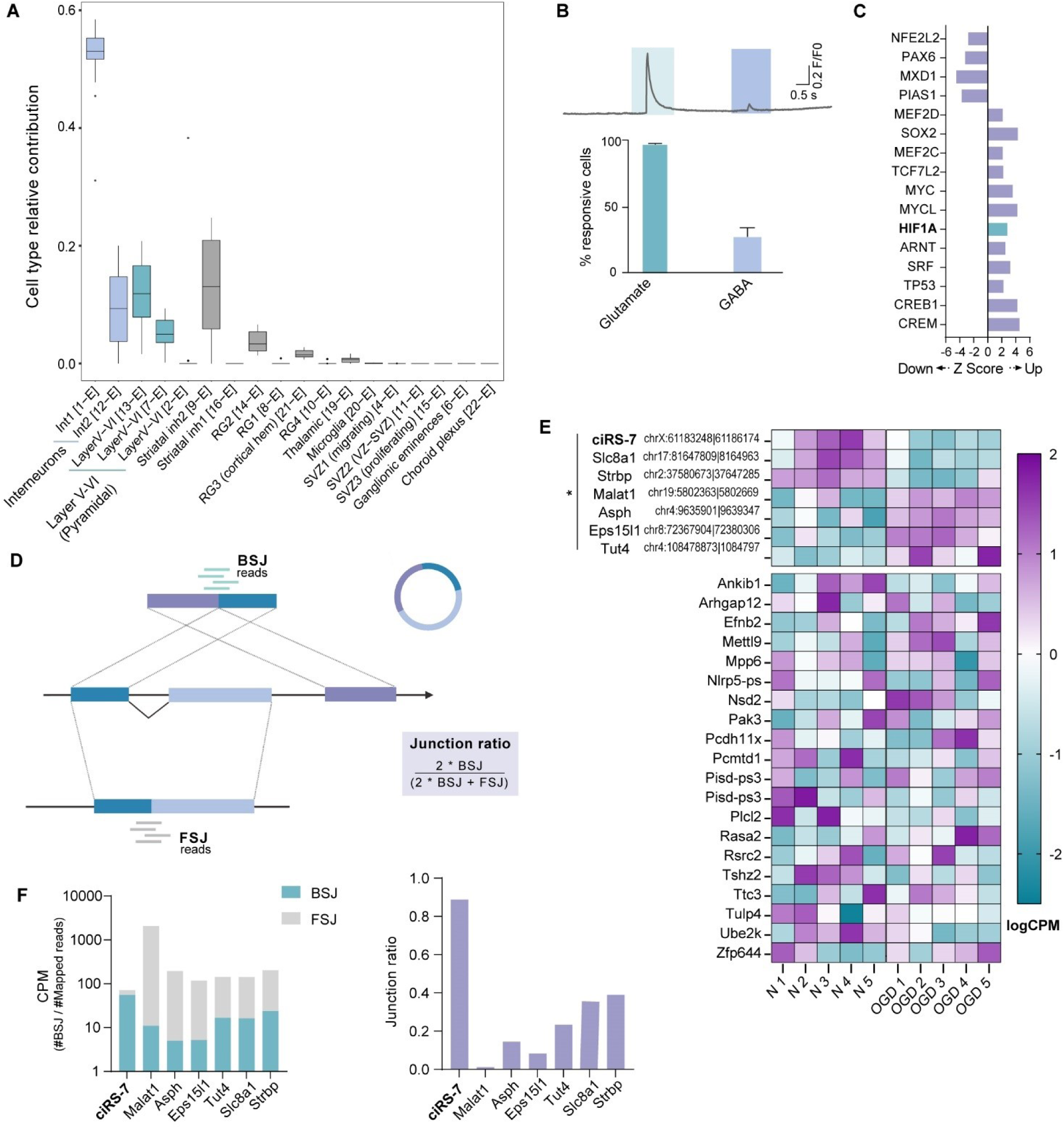
ciRS-7 is the most abundant circRNA in mouse cortical neuron cultures and downregulated by in vitro OGD. **(A)** Box plot of bulk RNA-seq deconvolution scores, indicating neuronal type contributions to overall gene expression profile. X-axis labels derive from initial categorization by the authors of the dataset (GSE123335), sorted by median of universal semantic groups (e.g., Int1, Int2 in Interneurons). **(B)** Calcium imaging analysis of cellular response to glutamate and GABA with representative trace (n = 11; signal > 5% of baseline; mean ± SD). **(C)** Bar plot of Ingenuity Pathway Analysis (IPA) of bulk RNA-seq data from OGD-treated cortical neurons. Z-score denotes prediction of transcription factor activation (positive) or inhibition (negative); significant HIF1α activation is highlighted in green (|z| > 2 is considered significant). **(D)** Schematic of CIRIquant algorithm: Gene with three exons (dark blue, light blue, purple) produces a circular molecule via back-splicing or a linear transcript. Green bars denotes specific circular reads on back-splice junction (BSJ), the reads from forward splicing junctions (FSJ) in the same region are in gray. The Junction ratio score formula calculates a circularization score, indicating the percentage of the gene in circular form. **(E)** Heat map of logCPM for detected circRNAs in normoxic (N) and oxygen/glucose-deprived (OGD) murine cortical neurons. Green indicates low expression, while purple indicates high expression. Significantly differentially expressed circRNAs are grouped with a bar (* = adj. p-value < 0.05, n = 5). **(F)** (*left*) Bar plot of CPM of the BSJ and FSJ of differentially expressed circularRNAs between normoxic and OGD conditions, BSJ represents circular specific reads. (*right*) Bar plot of the Junction ratio score bar plot from CIRIquant algorithm, showing the ratio of BSJ to FSJ reads mapped to the BSJ site and indicating the percentage of circularization relative to the linear transcript.

Ingenuity Pathway Analysis (IPA) of the differentially expressed genes between normoxic and OGD neurons confirmed the activation of Hypoxia-inducible factor 1-alpha (HIF-1alpha) (Figure 1C) and downstream upregulation of glycolysis, a characteristic hallmark of OGD^20^ (Supplementary Table S1, Supplementary Figure S1B). Moreover, colorimetric cell viability assay indicated 30% decrease in neuronal viability post-exposure (Supplementary Figure S1C), confirming vulnerability to ischemia-induced cell death.

CircRNAs, generated by back-splicing, are identified through back-splice junction reads (BSJ) spanning regions that are not present in regularly spliced transcripts (forward splice junction reads, FSJ) (Figure 1D). By using *CIRIquant* algorithm^21^, which comprises several circRNAs identification tools, we identified 27 circRNAs with high confidence, of which 7 were significantly differentially expressed (DE) between normoxic (control) and OGD conditions (Figure 1E, Supplementary Table S2). Notably, ciRS-7 was the most expressed circRNA yielding the highest number of back-splice junctions (Figure 1F, left) and the highest circular to linear ratio (Junction ratio) (Figure 1F, right). The obtained junction ratio score of above 0.88, indicate that the transcript generated by the ciRS-7 locus are over 88% in circular form, while other circRNAs exhibited only 40% of circular expression. Specifically, ciRS-7 was one of the three downregulated circRNAs upon OGD with a log_2_FC of - 0.552.

### ciRS-7 network is altered in conditions mimicking ischemic stroke in vitro

ciRS-7 is part of a feedback loop with miR-671 and miR-7 microRNA^5,7^ and indirectly, through miR-7, with the long non-coding RNA Cyrano^8^ (Figure 2A). To investigate the relative expression of these players in *in vitro* OGD conditions, we performed small RNA sequencing of cortical neurons exposed to OGD for 12h. Together with the previously presented dataset, we were able to capture circRNAs, mRNAs, long non-coding RNAs and small RNA transcripts (Supplementary Table S2).

**Figure 2.**
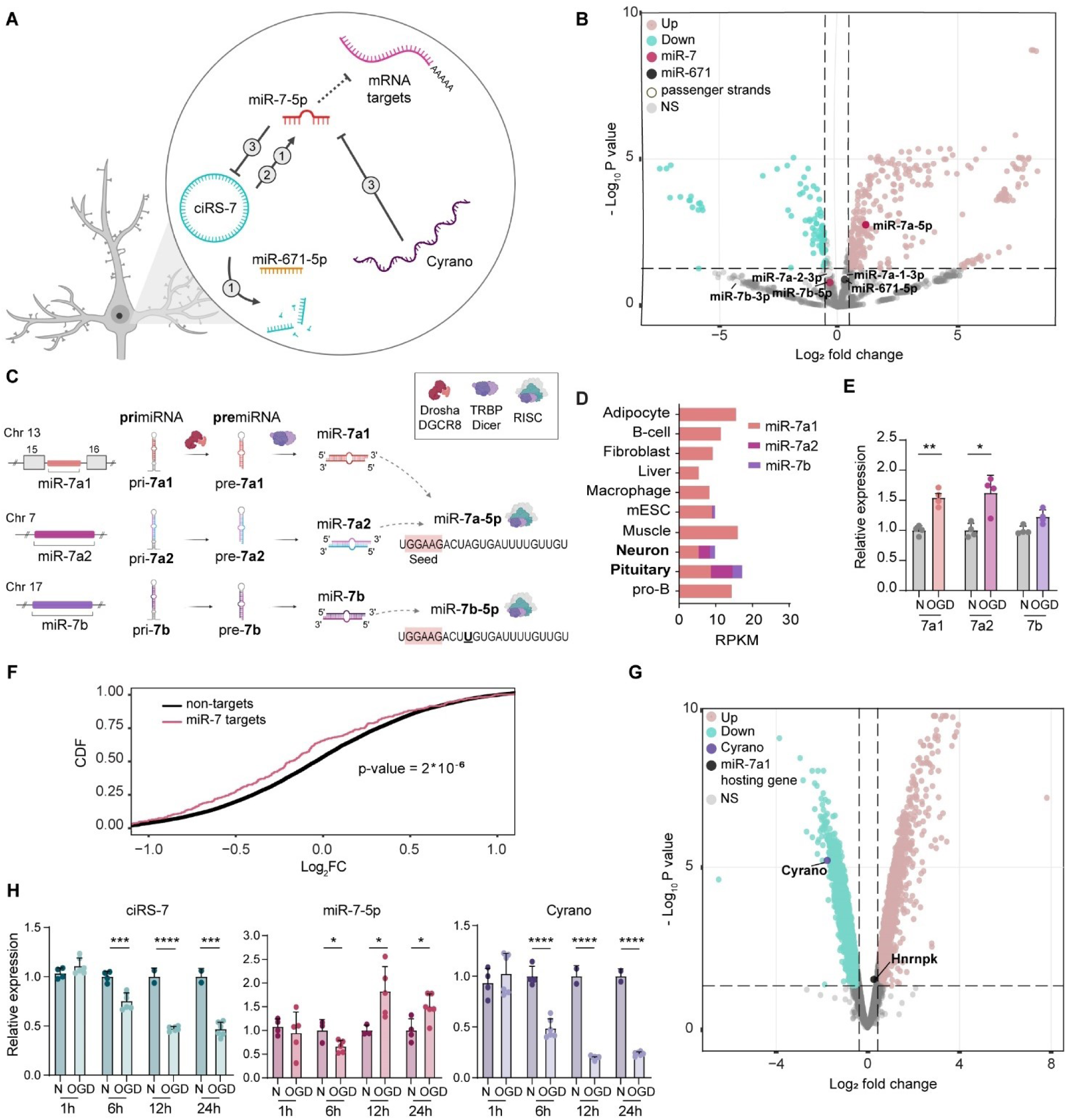
ciRS-7 network is dynamically altered in ischemic stroke like conditions. **(A)** Schematic of the regulatory network involving ciRS-7, miR-7, Cyrano, and miR-671. References: (1) Hansen et al., EMBO J, 2011; Hansen et al., Nature, 2013; (2) Piwecka et al., Science, 2017; (3) Kleaveland et al., Cell, 2018. **(B)** Volcano plot of differentially expressed microRNAs in OGD-treated cortical neuron cultures. Significantly upregulated (pink) and downregulated (green) microRNAs, miR-7 variants (red), miR-7 passenger strands (white), and miR-671 (black) are highlighted (n = 5; p-adj. < 0.05 and |log2FC| > 0.3). **(C)** Schematic of murine genomic loci illustrating the transcription of miR-7 primary transcripts (pri-miR), their processing into precursor molecules (pre-miR), and maturation into mature miR-7. The shared seed sequence between miR-7a-5p and miR-7b-5p is highlighted in red, and the non-seed mismatch at position 10 between the variants is underlined in black. **(D)** Bar plot of miR-7 loci reads per kilobase million (RPKM) in different cell type obtained from GRO-seq datasets. **(E)** Bar plot of miR-7 pri-miRNA expression in cortical neurons post-OGD treatment quantified by qPCR. Relative expression normalized to normoxic condition (n = 4; * p < 0.05, ** p < 0.01, paired t-test; mean ± SD).**(F)** Cumulative distribution functions (CDFs) plot of log fold changes of genes in OGD-treated wild-type cortical neurons (12h) compared to normoxic conditions. Red curve represents CDF for miR-7 targets, and black curve for non-targets (n = 5; p-value from Kolmogorov-Smirnov test). **(G)** Volcano plot of differentially expressed transcripts in OGD-treated cortical neuron cultures. Significantly upregulated (pink) and downregulated (green) transcripts, Cyrano (purple), and miR-7 hosting gene HnrnpK (black) are highlighted (n = 5; p-adj. < 0.05 and |log2FC| > 0.3). **(H)** Bar plot of ciRS-7 (blue), miR-7 (magenta), and Cyrano (purple) quantification by qPCR in OGD-treated cortical neurons at various timepoints (1h, 6h, 12h, and 24h). Relative expression normalized to normoxic condition (normoxia n = 2-4, OGD n = 5-6; * p < 0.05, *** p < 0.001, **** p < 0.0001, unpaired t-test; mean ± SD).

Small RNA-sequencing identified 333 upregulated and 88 downregulated miRNAs following OGD (Figure 2B). Notably, the levels of miR-671-5p remained unaltered (Figure 2B) which was confirmed independently by RT-qPCR (Supplementary Figure S2A), in contrast to previous reports of divergent miR-671-5p and ciRS-7 expression in different contexts^7–9^. Instead, we detected a significant upregulation in miR-7a-5p, but not of the variant miR-7b-5p (Figure 2B, Supplementary Table S2). Similarly to human, in mouse miR-7 is redundantly encoded by three different loci (*miR-7a-1*, *miR-7a-2*, *miR-7b*), each produced from different primary transcript (pri-miRNA) and precursor (pre-miRNA). Further processing, in all cases from −5p arm, give rise to two mature miR-7 sequences (miR-7a and miR-7b), differing only by a single nucleotide in the non-seed position number 10 (Figure 2C). To study the expression of the three miR-7 loci in our system, we utilized Global run-on sequencing (GRO-seq), which enable the capture of nuclear nascent RNA primary molecules, and compared the expression in our murine culture^22^ with different mouse tissues and cell lines^22–32^ (Supplementary Table S3). In physiological conditions, only cortical neurons and the pituitary gland actively transcribe all three independently regulated *miR-7* loci (Figure 2D, Supplementary figure S3). The active transcription from three different loci in neuronal cells suggests a higher order of regulation of this miRNA which influences the mature forms. Under OGD conditions, only the pri-miRNAs contributing to the expression of miR-7a (miR-7a-1 and miR-7a-2) were significantly upregulated (Figure 2E), indicating that the observed upregulation occurs already at the level of transcription. None of miR-7 passenger strands generated from the three precursors (miR-7a-1-3p, miR-7a-2-3p, and miR-7b-3p) were altered in our sequencing data (Supplementary Table S2). Moreover, we detected no changes in the host gene in which miR-7a-1 is embedded *(Hnrnpk*), and from which the most abundant miR-7 primary transcript is generated (Supplementary Table S2). Taken together these data show that OGD specifically regulates miR-7a-5p variant at the transcriptional and/or post-transcriptional level.

To test the possible functional relevance of the upregulation of miR-7a-5p, we acquired predicted and validated targets of miR-7a-5p using *miRWalk* algorithm analysis^33,34^ and compared their overall expression in conditions of OGD. In accordance with the upregulation of miR-7, miR-7 targets showed a significant overall downregulation (Figure 2F, Supplementary Table S4, Supplementary Figure S2B), suggesting a canonical functional repressive role of this miRNA.

The role of miR-7 interaction with ciRS-7 remains controversial, and it is thought to set the balance between a positive and negative feedback^8,9^ (Figure 2A). Additionally, Cyrano promotes target-directed miRNA degradation (TDMD) of miR-7 through a site of almost perfect complementarity, indirectly regulating ciRS-7 levels and localization^8^ (Figure 2A). In our dataset the lncRNA Cyrano was significantly downregulated upon OGD (Figure 2G), in line with the observed upregulation in miR-7a-5p (Figure 2B). To identify a first responder to OGD in the context of this molecular circuitry we evaluate the time-dependent changes in the levels of these molecules. We subjected cortical neurons to OGD for 1, 6, 12 and 24 hours and assessed the expression level of ciRS-7, miR-7 and Cyrano. We detected a significant downregulation of ciRS-7 and Cyrano already at 6 hours after OGD, prior to the upregulation of miR-7 at 12 hours (Figure 2H). Interestingly, our data revealed a significant downregulation of miR-7 at 6h of OGD concomitantly with downregulation in ciRS-7 and Cyrano, which is in line with the previously suggested role of ciRS-7 in stabilizing miR-7^9^.

### ciRS-7 KO neurons exhibit a distinct outcome after OGD without altering the overall OGD response

The absence of ciRS-7 alters synaptic transmission in excitatory neurons and produces schizophrenia-like phenotype *in vivo*^9^, connecting this molecule to glutamatergic transmission. This is of particular interest in conditions of ischemic stroke, where glutamate mediated excitotoxicity is a critical contributor to neuronal cell death^12^. Our cortical neuron culture enriched in glutamatergic neurons (Figure 1B) provides a system in which these events can be studied *in vitro.* Neurons cultured from ciRS-7 KO mice showed higher Ca^2+^-responses to the stimulation with glutamate compared to their wild-type (WT) counterparts (Figure 3A). Consequently, ciRS-7 KO neurons also exhibited a significantly higher sensitivity to excitotoxicity upon exposure to high concentration of glutamate (Figure 3B).

**Figure 3.**
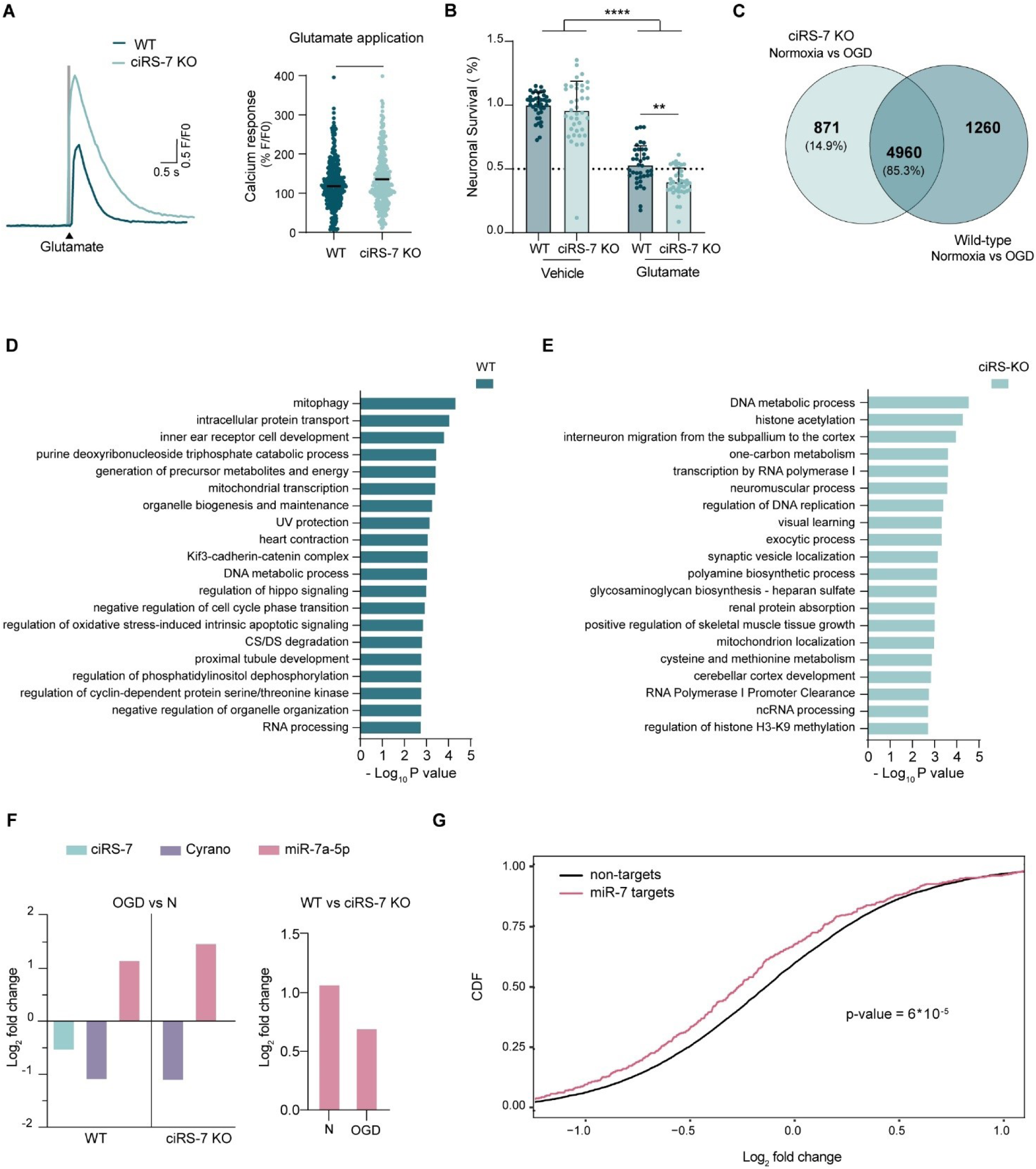
ciRS-7 KO genotype exhibits differential OGD phenotypical output without affecting the OGD induced changes in miR-7 and Cyrano. **(A)** (Left) Representative trace of calcium-induced fluorescence in response to glutamate treatment in wild-type and ciRS-7 KO cortical neurons. (Right) Scatter dot plot of calcium imaging analysis showing the cellular response to glutamate in wild-type and ciRS-7 KO cortical neurons (n = 8; * p < 0.05, unpaired t-test; central bar represents median value). **(B)** Bar plot of relative absorbance in MTT viability assay for wild-type and ciRS-7 KO cortical neurons treated with vehicle or 250μM glutamate. Data presented as survival % normalized to wild-type vehicle (n = 4; * p < 0.05, **** p < 0.0001, one-way ANOVA corrected with Tukey’s post-hoc test; mean ± SD). **(C)** Venn diagram of differentially expressed genes in normoxic versus OGD conditions of wild-type and ciRS-7 KO cortical neurons (n = 5; p-adj. < 0.05 and |log2FC| > 0.3). **(D)** Bar plot of functional enrichment analysis top 20 significant Metascape clusters performed on differentially expressed genes in wild-type normoxic versus OGD cortical neurons. **(E)** Bar plot of top 20 significant Metascape clusters from functional enrichment analysis on differentially expressed genes in wild-type normoxic versus OGD cortical neurons. **(F)** (Left) Bar plot of log2FC for ciRS-7 (blue), miR-7 (magenta), and Cyrano (purple) from RNA-seq of wild-type and ciRS-7 KO cortical neurons in normoxic conditions versus after 12h of OGD treatment. (Right) Bar plot of log2FC for miR-7 (magenta) from RNA-seq, highlighting expression differences in normoxic conditions (N) or OGD between wild-type and ciRS-7 KO genotypes (n = 5; p-adj. < 0.05 and |log2FC| > 0.3). **(G)** Cumulative distribution functions (CDFs) plot of log fold changes of genes in OGD-treated ciRS-7 KO cortical neurons (12h) compared to normoxic conditions. The red curve represents CDF for miR-7 targets, while the black curve represents non-targets (n = 5; p-value from Kolmogorov-Smirnov test).

To test if this increased sensitivity to glutamate is affecting gene expression changes in OGD, we subjected the ciRS-7 KO cortical cultures to OGD for 12 hours, following the previous experimental design (Supplementary Figure S1A) and performed total RNA-seq. OGD induced deregulation in total of 5767 genes in ciRS-7 KO neurons compared to normoxic conditions, of which the 85% were shared with WT neurons (Figure 3C, Supplementary Table S5). Consistent with this, the analysis of interaction terms in the differential expression analysis, aimed at distinguishing differences in the OGD response between WT and ciRS-7 KO, revealed no significant difference in the overall OGD response between the two genotypes (Supplementary Table S6). However, 871 genes were differentially expressed only in ciRS-7 KO neurons, while the expression of 1260 genes was exclusively altered in WT neurons subjected to OGD. ciRS-7 WT specific genes altered in OGD were functionally enriched in mitochondrial processes, DNA metabolism, regulation of cell-cycle and oxidative stress-induced apoptosis (Figure 3D). The 871 KO-specific genes were instead involved in developmental and pro-regenerative processes (*progenitor migration to the cortex, DNA replication, transcription, cerebellar cortex development*) (Figure 3E). Despite the genetic predisposition to glutamate sensitivity of the ciRS-7 KO, which would suggest a deleterious outcome in the response to OGD, ciRS-7 KO neurons regulate pathways of resilience during OGD. In support of this, ciRS-7 KO neurons did not show increased cell death compared to WT in response of OGD treatment (Supplementary Figure S2C). This result implies a distinction in OGD outcome not attributed to a change in the OGD response itself, but rather to the regulation of the cascade of processes downstream of the response.

Interestingly, the lack of ciRS-7 did not affect OGD induced changes in the expression of miR-7 and Cyrano (Figure 3F). Similar to WT neurons, Cyrano lncRNA was downregulated in ciRS-7 KO neurons responding to OGD (Figure 3F, Supplementary Table S5). Moreover, even though ciRS-7 KO neurons stably express lower levels of miR-7a-5p^9^ (Figure 3F, Supplementary Table S5), small-RNA sequencing revealed a significant upregulation of miR-7a-5p in response to OGD also in these neurons (Figure 3F, Supplementary Table S5). In line with our previous findings, we detected no changes in miR-671-5p, miR-7b-5p and miR-7 passenger strand expression (Supplementary Table S5). Nonetheless, whilst the expression in miR-7a-5p in OGD remained lower in ciRS-7 KO neurons compared to WT neurons (Figure 3F), the targets of miR-7a-5p, both predicted and validated, showed a significant global repression in these conditions (Figure 3G).

### Alteration of ciRS-7 affects ischemic stroke outcome in vivo

To assess the reproducibility of our findings in ischemic stroke *in vivo*, *BALB/c* mice were subjected to permanent middle cerebral artery occlusion (pMCAO) and levels of ciRS-7, Cyrano and miR-7a-5p were evaluated by qPCR at six hours, one day and five days post-ischemia. In line with our *in vitro* results, we detected a significant downregulation of ciRS-7 and Cyrano and upregulation of miR-7a-5p in the peri-ischemic cortex (PI) at one day post-ischemia (dpi) compared to contralateral cortex (CL) (Figure 4A). Moreover, similar to OGD, we failed to detect alterations in miR-671-5p levels (Supplementary Figure S2D).

**Figure 4.**
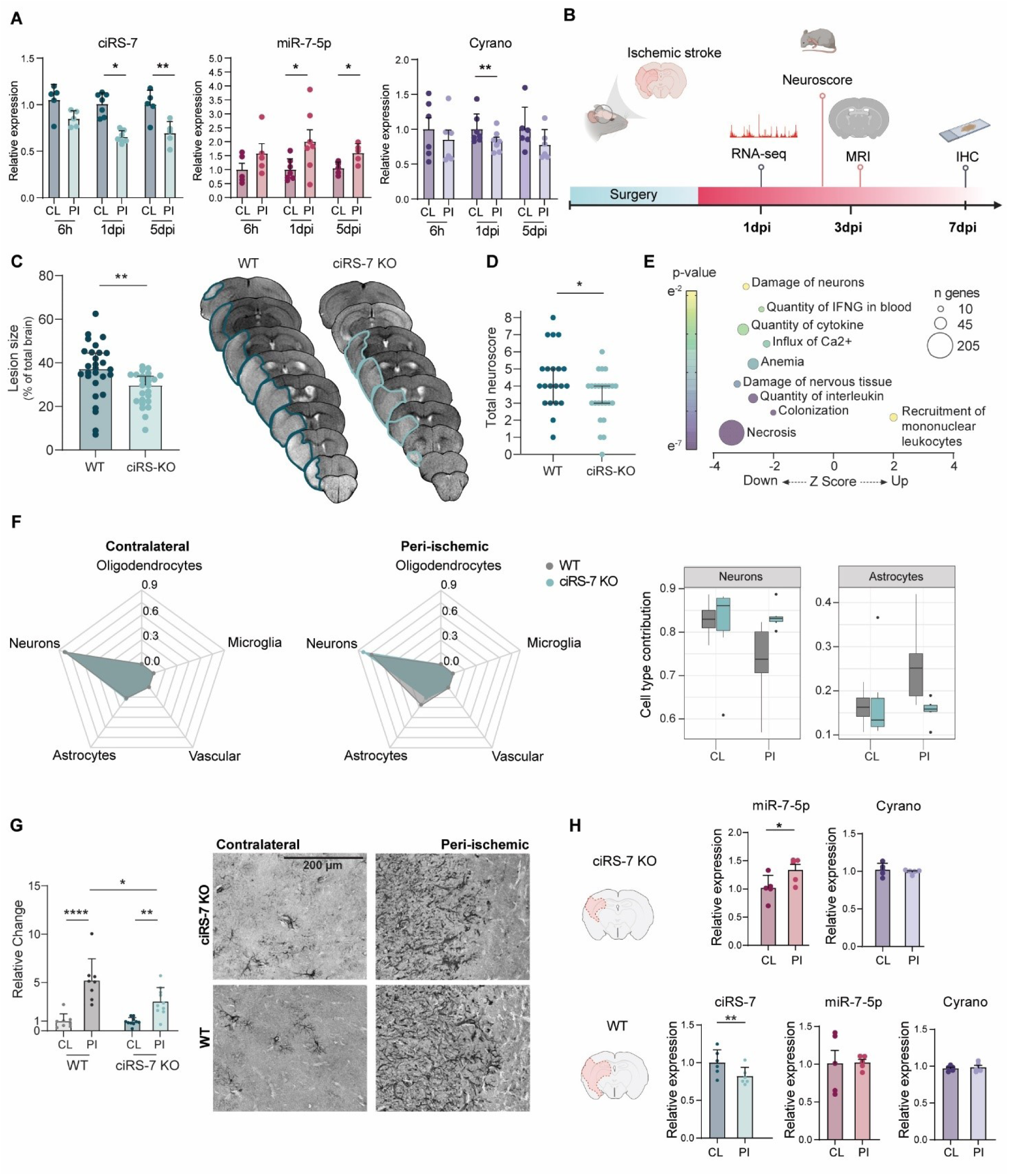
Lack of ciRS-7 ameliorates ischemic stroke outcome in vivo. **(A)** Bar plot of qPCR quantification for ciRS-7 (blue), miR-7 (magenta), and Cyrano (purple) in 3-4 months old BALB/c mice after pMCAo surgery. Peri-ischemic (PI) and contralateral (CL) cortices were collected at different timepoints (6h, 1d, and 5d). Relative expression normalized to the average CL expression (n = 6-7; * p < 0.05, *** p < 0.001, **** p < 0.0001, paired t-test; mean ± SD). **(B)** Schematic of the experimental design: Wild-type and ciRS-7 KO mice underwent tMCAO surgery, followed by neuroscore testing and MRI monitoring at 1, 3, and 7 days post-surgery (dpi). At 1dpi, six mice were utilized to obtain peri-ischemic and contralateral cortices for RNA-sequencing. Immunohistochemistry analysis was performed at later timepoints using eight to nine mice per group. **(C)** (Left) Bar plot of MRI quantification of the lesion size in wild-type and ciRS-7 KO mice 1 day post-tMCAO surgery. Data presented as lesion percentage on total brain size adjusted for edema (n = 25-28 per group; ** p < 0.01, Mann–Whitney test; median with 95% confidence interval). (Right) Representative MRI image illustrating changes in lesion perimeter (blue) between wild-type (WT) and ciRS-7 KO animals. **(D)** Scatter dot plot of sensorimotor deficits assessed by neuroscore in tMCAO wild-type and ciRS-7 KO mice at 1dpi. Data presented as total neuroscore (n = 22-25 per group; * p < 0.05, Mann–Whitney test; median with 95% confidence interval). **(E)** Bubble plot of Ingenuity Pathway Analysis (IPA) for differentially expressed genes between wild-type and ciRS-7 KO peri-ischemic cortices from RNA-seq. Z-score reflects IPA prediction of pathway activation (positive) or inhibition (negative) in ciRS-7 KO. Color bar indicates p-value significance (yellow to purple), and bubble size represents the number of genes in the pathway (|z| > 2 considered significant; pathways with >10 genes included). **(F)** (Left) Spiderweb plot of bulk RNA-seq deconvolution scores of wild-type (gray) and ciRS-7 KO (blue) contralateral and peri-ischemic cortices. (Right) Bar plot illustrating neurons and astrocyte contributions based on deconvolution scores in wild-type (gray) and ciRS-7 KO (blue) contralateral and peri-ischemic regions. **(G)** (Left) Bar plot of quantification for GFAP DAB (3,3′-Diaminobenzidine) immunostaining in wild-type and ciRS-7 KO tMCAO mice at 7dpi. (Right) Representative images of the staining highlighting GFAP^+^ cells. Relative expression normalized to the average contralateral expression of wild-type mice (n = 8; * p < 0.05, ** p < 0.01, **** p < 0.0001, one-way ANOVA corrected with Tukey’s post-hoc test; mean ± SD). **(H)** Bar plot of qPCR quantification for ciRS-7 (blue), miR-7 (magenta), and Cyrano (purple) in contralateral (CL) and peri-ischemic (PI) cortices of ischemic wild-type and ciRS-7 KO mice at 1dpi. Schematic representation correlates with MRI lesion size at the same timepoint. Data normalized to the average contralateral expression of wild-type mice (n = 5 per group; * p < 0.05, ** p < 0.01, paired t-test; mean ± SD).

To explore the functional implications of ciRS-7 KO neurons ambiguous behavior involving glutamate sensitivity without increased cell death during OGD, we tested whether ciRS-7 KO mice exposed to transient ischemic stroke would show altered vulnerability to ischemic damage and associated sensorimotor impairments. The ischemic lesion size was measured using Magnetic Resonance Imaging (MRI) and behavioral deficits were assessed using Neurological Severity Scores (NSS) at acute (1dpi), subacute (3dpi) and chronic (7dpi) timepoints after transient middle cerebral artery occlusion (tMCAO) (Figure 4B). At 1dpi, ciRS-7 KO mice showed a significant reduction in the ischemic lesion volume compared to their WT controls (Figure 4C) and significantly ameliorated motor deficits (Figure 4D). Interestingly, ciRS-7 KO animals lesion size changes occurred transiently at the acute timepoint (1dpi), without delaying damage progression (Supplementary Figure S4A,B). To identify the molecular changes associated with acute reduction in the lesion size and motor deficits in ciRS-7 KO mice, mice were sacrificed at 1dpi and peri-ischemic and the contralateral cortex was used for mRNA sequencing. IPA analysis of differentially expressed genes in the peri-ischemic cortex of ciRS-7 KO animals indicated substantial inhibition in pathways related to neuronal and tissue damage, cytokine and interleukin release, and calcium influx compared to WT controls (Figure 4E, Supplementary Table S7). Moreover, most differentially expressed genes were mostly contributing to pathways of *necrosis* (205 genes) and *quantity of cytokine* (41 genes), suggesting that the decreased lesion size in the ciRS-7 KO mice may be due to inhibition of necrotic pathways and reduction in cytokine release.

Aiming to identify crucial cellular responders mediating the observed gene expression changes, we carried out deconvolution analysis by using adult mouse brain scRNA-seq data^35^ to assess the contribution of different cell populations in our bulk mRNA sequencing dataset. This analysis identified neurons and astrocytes as major cellular responders contributing to WT and ciRS-7 KO gene expression profiles (Figure 4F). In line with the significantly reduced tissue death measured by MRI (Figure 4C), ciRS-7 KO mice exhibited increased neuronal involvement in the peri-infarct cortex compared to WT mice, accompanied by a decrease in astrocyte contribution. To confirm this, brain slices from ischemic ciRS-7 KO mice were collected at the peak of the immune response (7dpi) ^36^ and stained for astrocytic glial fibrillary acidic protein (GFAP). Although ischemic stroke induced GFAP expression in the peri-infarct cortex for both WT and ciRS-7 KO mice, ciRS-7 KO mice exhibited significantly reduced astrogliosis compared to WT animals (Figure 4G).

At molecular level, tMCAO significantly downregulated ciRS-7 in WT mice at 1dpi in the peri-ischemic cortex, while miR-7-5p and Cyrano levels remained unaltered (Figure 4H, Supplementary Figure S4C). However, ciRS-7 KO mice showed a significant upregulation of miR-7-5p and unaltered Cyrano levels (Figure 4H). These data suggest a correlation between the absence of ciRS-7 and a faster upregulation of miR-7 in response to transient ischemic stroke and a marginal role of Cyrano in this system at this timepoint.

### ciRS-7 prevents miR-7 effects on dampening glutamatergic signaling in excitatory neurons

Previous studies linked miR-7 to oxidative stress response^37^, in particular in low glucose conditions^38^. To establish the possible functional link between the upregulation of miR-7 in ciRS-7 KO mice in conditions of ischemic stroke, we subjected cre-loxP inducible miR-7 KO mice^39^ to tMCAO (Figure 5A, Supplementary Figure S5A,B). At the acute timepoint (1dpi), no changes in lesion volume, as measured by MRI, were detected between miR-7 KO and WT mice (Supplementary Figure S5C). Accordingly, miR-7 KO mice did not show any differences in motor deficits compared to their WT controls (Supplementary Figure S5D). However, miR-7 KO mice (Figure 5B) exhibited an increase in the lesion volume compared to their WT controls at later timepoint (7 dpi), unrelated to changes in Neuroscore (Supplementary Figure S5E), indicating a potential role of miR-7 in regulating ischemia-induced secondary cell death.

**Figure 5.**
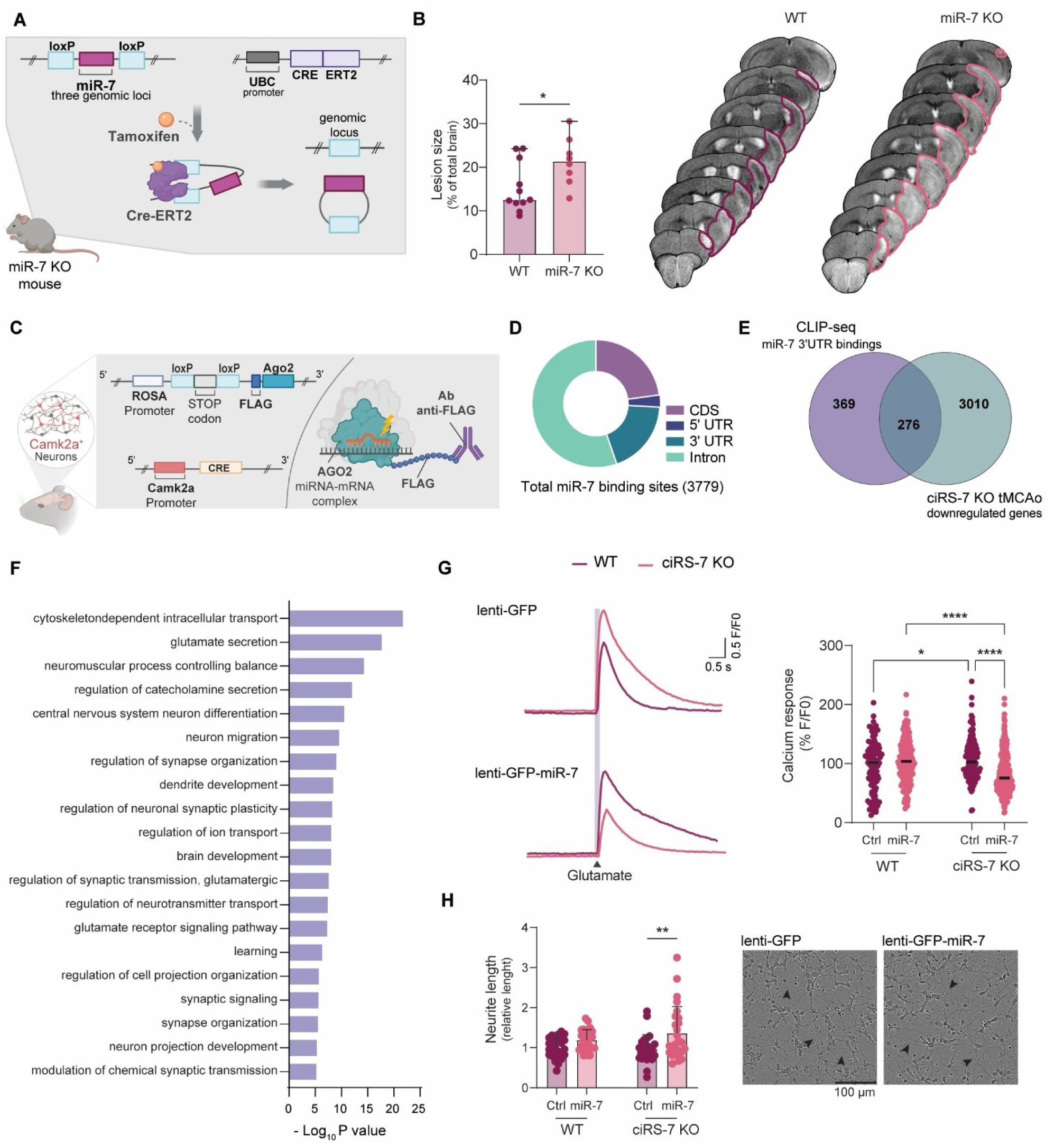
Lack of miR-7 exaggerates ischemic stroke outcome regulating glutamatergic response. **(A)** Schematic of Cre-LoxP miR-7 KO animal model: mice have a transgenic genome with LoxP sequences (blue) flanking all three miR-7 loci (magenta), and tamoxifen (orange)-inducible Cre recombinase (purple). Tamoxifen activation of Cre recombinase leads to miR-7 loci deletion. Control mice lack the Cre recombinase transgene. **(B)** (Left) Bar plot of MRI quantification for lesion size in wild-type and miR-7 KO mice one day post-tMCAO surgery. Data presented as lesion percentage on total brain size adjusted for edema (n = 8-11 per group; * p < 0.05, Mann–Whitney test; median with 95% confidence interval). (Right) Representative MRI image illustrating changes in lesion perimeter (blue) between wild-type (WT) and miR-7 KO animals. **(C)** Schematic of recombinant mice for generating Ago2 CLIP-seq data in excitatory neurons: FLAG-Ago2 (blue) is selectively translated in Cam2Ka^+^ (red) excitatory neurons, where Cre recombinase (yellow) expression removes a stop codon (gray) leading to FLAG-Ago2 translation. Following cross-linking (thunder), the FLAG-Ago2-miRNA-target complex is immunoprecipitated with an anti-FLAG antibody (purple), and the sequenced output represents the miRNA-target complex. **(D)** Pie chart illustrating the distribution of miR-7 binding sites from CLIP-seq analysis. Among the 3779 identified binding sites, categories include protein coding sequences (CDS, purple), 5’ untranslated regions (5’ UTR, blue), 3’ untranslated regions (3’ UTR, dark green), and intronic sequences (Intron, light green). **(E)** Venn diagram depicting the miR-7 binding sites in the 3’ UTR identified by CLIP-seq in excitatory neurons (645) and the downregulated genes in the peri-ischemic region of ciRS-7 KO tMCAO animals (3286), where miR-7 is upregulated. The overlap (276 genes) represents potential miR-7 physical targets affected in ischemic stroke. **(F)** Bar plot of the top 20 significant Metascape clusters from functional enrichment analysis on genes identified as miR-7 physical targets potentially affected in ischemic stroke. **(G)** (Left) Representative trace of calcium-induced fluorescence in response to glutamate treatment in wild-type and ciRS-7 KO neurons infected with miR-7 or control lentivirus. (Right) Scatter dot plot of calcium imaging analysis quantifying the cellular response to glutamate in wild-type and ciRS-7 KO cortical neurons infected with control (GFP) or miR-7 overexpressing (GFP + miR-7) virus (n = 5; * p < 0.05, **** p < 0.0001, Kruskal–Wallis test; central bar represents median value). **(H)** (Left) Bar plot of neurite length quantified through live imaging of wild-type and ciRS-7 KO cortical neurons infected with control (GFP) or miR-7 overexpressing lentivirus (GFP + miR-7). Data normalized to the average neurite length of wild-type GFP-infected neurons (n = 3 biological replicates, n = 8 technical replicates; ** p < 0.01, one-way ANOVA test corrected with Tukey’s post-hoc test; mean ± SD). (Right) Representative image of ciRS-7 KO cortical neurons infected with control (GFP) or miR-7 overexpressing virus (GFP + miR-7).

To understand the functional role of miR-7 in ischemic stroke, we first validated miR-7 functional target sites in our system by analyzing miR-7 target binding sites in an Ago2 HITS-CLIP sequencing dataset of pyramidal excitatory neurons^40^ (Figure 5C, Supplementary Table S8). By identifying RISC-associated binding sites on expressed transcripts in mature pyramidal neurons, our analysis of the minimal seed site hexamer of miR-7 revealed 645 potential targets in the 3’ UTRs (Figure 5D). As expected, ciRS-7 had the highest number of functional binding sites (n=139), representing over 11% of the total reads (Supplementary Table S8). The intersection of Ago2 HITS-CLIP physical targets and the downregulated genes in the peri-ischemic cortex of ciRS-7 KO mice, where miR-7 is upregulated, highlighted 276 common genes (Figure 5E). These genes were functionally enriched in pathways related to glutamatergic synaptic transmission and morphological changes of neuronal projections (Figure 5F, Supplementary Figure S6).

To discern the specific contributions of ciRS-7 and miR-7 to glutamatergic signaling and neuronal morphological changes, we opted to replicate the molecular changes associated with low ciRS-7 and high miR-7 *in vitro.* We transduced ciRS-7 WT and KO neurons with lentivirus overexpressing GFP and miR-7a or GFP only, under the neuronal specific human Synapsin 1 (hSYN) promoter. Infection efficiency was evaluated through fluorescence microscopy (Supplementary Figure S7A) and we confirmed by RT-qPCR that miR-7a-5p levels resembled the endogenous levels under OGD condition (Supplementary Figure S7B). Whilst overexpression of miR-7 in WT neurons did not affect the overall neuronal glutamate excitability, increased levels of miR-7 in ciRS-7 KO neurons significantly reduced their response to glutamate (Figure 5G), thus reverting the genotype of glutamate sensibility of the ciRS-7 KO neurons. This appeared to be a glutamate specific effect, as the overexpression of miR-7 did not affect GABAergic response (Supplementary Figure S7C).

Finally, to evaluate the functional impact of the potential targets of miR-7 involved in morphological changes of neuronal projections (Figure 5F), we performed live imaging analysis of neurite outgrowth of ciRS-7 KO and WT neurons infected with miR-7 lentivirus. In agreement with the calcium imaging experiment, we were able to detect a significant increase in the neurite length when over expressing miR-7 in ciRS-7 KO cells, but not in WT neurons (Figure 5H). This aligns with a previous study showing a direct impact of miR-7 overexpression on neurite length in a neuroblastoma cell line^41^. Considering circRNAs are typically low in cell lines due to their rapid proliferation^42^, we questioned whether the effect of miR-7 on neurite length was due to the absence of ciRS-7 interference.

However, circRNAs are known to be poorly expressed in cell lines due of their high proliferation rate^42^. So, we wondered whether this miR-7 direct effect on neurite length was due to lack of ciRS-7 interference. We confirmed that the murine neuroblastoma cell line N2A expresses negligible ciRS-7 levels even when differentiated with retinoid acid, despite higher levels of Cyrano and miR-7 (Supplementary Figure S8A,B,C).

### miR-7 modulates AMPA-mediated glutamatergic signaling via ciRS-7/Cyrano network in ischemic conditions

To further unravel the intricate interactions within the miR-7/ciRS-7/Cyrano network, we conducted gain and loss of function experiments to modulate miR-7 expression in the context of ischemic stroke. We utilized a lentivirus expressing the Cyrano CR1 region, known for its role in mediating miR-7 degradation through TDMD^8^. Wild-type cortical neurons were transduced with lentivirus carrying CR1 or a control region featuring a mutated miR-7 binding site before exposure to a 12-hour period of OGD. As expected, overexpression of Cyrano CR1 region led to a decrease in miR-7 expression in normoxic conditions. Additionally, it prevented the OGD-induced increase in miR-7, in stark contrast to the mutated CR1, which had no impact on OGD-induced upregulation of miR-7 (Fig. 6A). Remarkably, preventing the increase in miR-7 expression preserved the levels of ciRS-7 during OGD, disclosing a previously proposed antagonistic relationship between these molecules^8^. The overexpression of the synthetic construct had no influence on the OGD-induced downregulation of the endogenous full-length Cyrano. Conversely, lentiviral induced overexpression of miR-7 induced an amplified miR-7 response during OGD (Fig. 6B). This molecular phenotype was consistently associated with a downregulation of ciRS-7, even under physiological conditions, further suggesting a potential involvement of miR-7 expression in the downregulation of ciRS-7. As expected, miR-7 overexpression had no effect on Cyrano basal expression or OGD-induced Cyrano downregulation (Fig. 6B).

**Figure 6.**
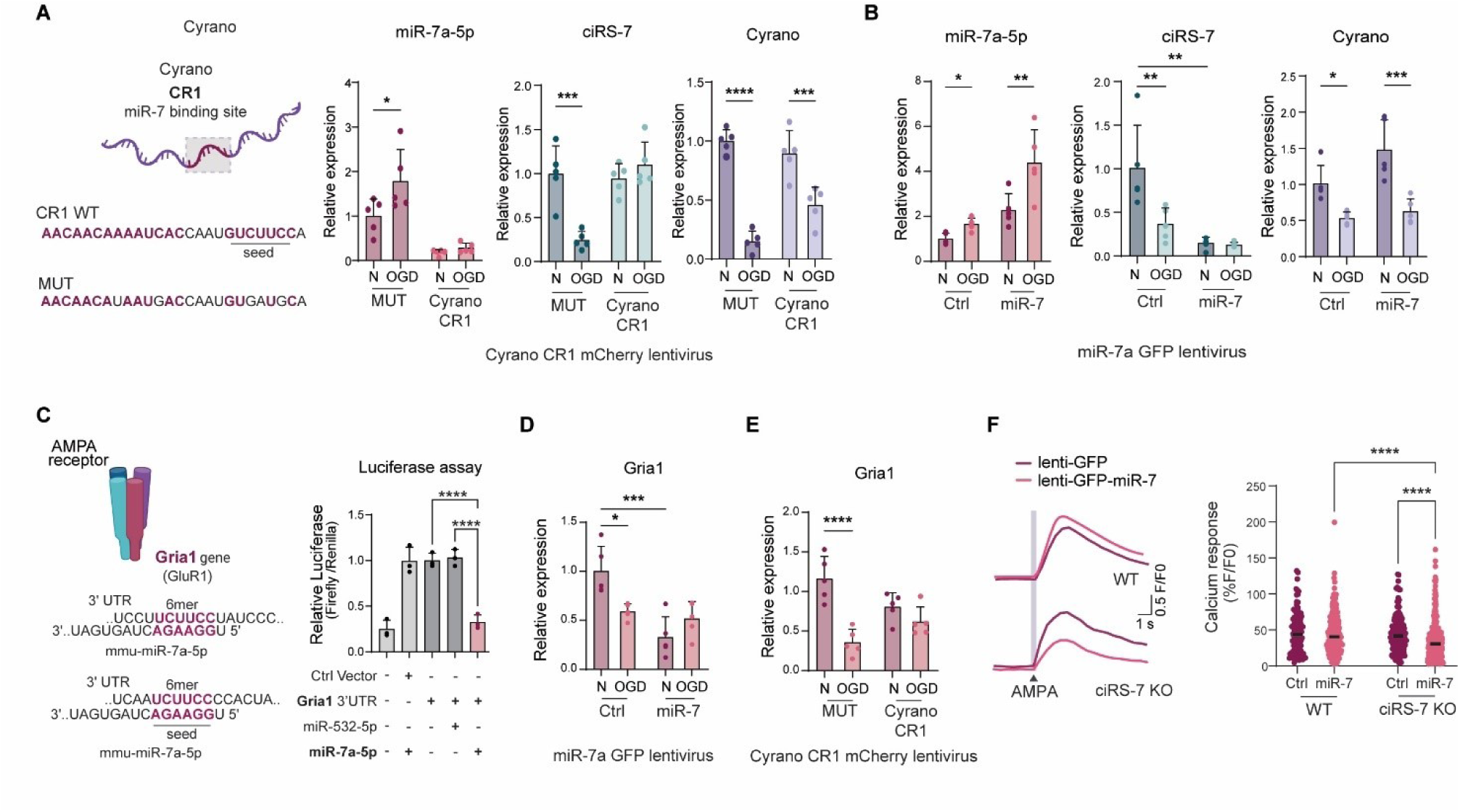
miR-7 regulation in OGD depends on ciRS-7 and Cyrano and modulate AMPA response. **(A)** (Left) Schematic of the wild-type Cyrano CR1 region (CR1) and its miR-7 binding sites mutated version (MUT). Magenta indicates pairing with miR-7a-5p, encompassing both seed region and a more extensive one. (Right) Bar plot of qPCR quantification for miR-7 (magenta), ciRS-7 (blue), and Cyrano (purple) in cortical neurons overexpressing Cyrano CR1 region (Cyrano CR1) or its mutated version (MUT) along with mCherry, treated with OGD for 12 hours. Relative expression normalized to normoxic condition of the mutated lentivirus (n = 5; * p < 0.05, *** p < 0.001, **** p < 0.0001, one-way ANOVA corrected with Tukey’s post-hoc test; mean ± SD). **(B)** Bar plot of qPCR quantification for miR-7 (magenta), ciRS-7 (blue), and Cyrano (purple) in cortical neurons overexpressing miR-7 GFP or a control vector with only GFP (Ctrl), treated with OGD for 12 hours. Relative expression normalized to normoxic condition of the control lentivirus (n = 5; * p < 0.05, ** p < 0.01, *** p < 0.001, one-way ANOVA corrected with Tukey’s post-hoc test; mean ± SD). **(C)** (Left) Schematic of the heteromeric AMPA receptor with its four subunits highlighted in different colors. Gria1, encoding the GluR1 subunit, is highlighted in magenta. Two 6mer sites on Gria1 3’ UTR and their potential pairing with miR-7a-5p in the seed region are shown (underlined). (Right) Bar plot of luciferase assay fluorescence quantification in HEKT293 cells transfected with control vector (Ctrl vector), Gria1 3’ UTR, and miR-532-5p (lacking binding sites) or miR-7a-5p (magenta). Relative expression normalized to Gria1 3’ UTR vector alone (n = 3; **** p < 0.0001, one-way ANOVA corrected with Tukey’s post-hoc test; mean ± SD). **(D)** Bar plot of qPCR quantification for Gria1 (magenta) in cortical neurons overexpressing miR-7 GFP or a control vector with only GFP (Ctrl), treated with OGD for 12 hours. Relative expression normalized to normoxic condition of the control lentivirus (n = 5; * p < 0.05, *** p < 0.001, one-way ANOVA corrected with Tukey’s post-hoc test; mean ± SD). **(E)** Bar plot of qPCR quantification for Gria1 (magenta) in cortical neurons overexpressing Cyrano CR1 region (Cyrano CR1) or its mutated version (MUT) along with mCherry, treated with OGD for 12 hours. Relative expression normalized to normoxic condition of the mutated lentivirus (n = 5; **** p < 0.0001, one-way ANOVA corrected with Tukey’s post-hoc test; mean ± SD). **(F)** (Left) Representative trace of calcium-induced fluorescence in response to AMPA treatment in wild-type and ciRS-7 KO cortical neurons transduced with miR-7 or control lentivirus. (Right) Scatter dot plot of calcium imaging analysis of the cellular response to AMPA in wild-type and ciRS-7 KO cortical neurons infected with control (GFP) or miR-7 overexpressing (GFP + miR-7) virus (n = 6-7; **** p < 0.0001, Kruskal–Wallis test; central bar represents median value).

By leveraging miR-7 targets validated through CLIP-seq that were dysregulated in ischemic mice (Fig. 5E, Supplementary Table S7), we pinpointed Gria1, a gene encoding a subunit of the AMPA receptor, critical for mediating glutamate excitotoxicity in ischemic stroke pathophysiology^43^. The 3’ UTR of Gria1 contains two potential 6mer binding sites for miR-7a-5p (Fig. 6C). To validate the physical interaction of these binding sites with miR-7a, we co-expressed a luciferase vector containing the 3’ UTR of Gria1 and miR-7a-5p in the HEK293T cell line. We observed a significant post-transcriptional canonical effect of miR-7 on the 3’UTR which was absent with an unrelated control miRNA (Fig. 6C). To further confirm the significance of this relationship in conditions of ischemic stroke, we measured the levels of Gria1 in cortical neurons transduced with miR-7 lentivirus prior OGD for 12h. In line with the RNA-seq data (Supplementary Figure S2B), OGD treatment led to a decrease in Gria1 expression in neurons transduced with the control vector. However, overexpression of miR-7 resulted in a 50% reduction in Gria1 mRNA levels under normal oxygen conditions, with no further alterations during OGD (Fig 6D). Overexpression of Cyrano CR1 region in neurons subjected to OGD prevented the OGD-induced downregulation of Gria1 (Fig. 6E). To demonstrate that the miR-7-mediated effect on Gria1 lead to a decrease response to AMPA, we carried out calcium imaging in ciRS-7 KO neurons that were lentivirally transduced to overexpress miR-7. In these conditions, ciRS-7 buffering effect on miR-7 is disrupted. As expected, the upregulation of miR-7 had no impact on wild-type neurons. However, in ciRS-7 KO neurons, we observed a significant decrease in AMPA-induced Ca^2+^ responses in miR-7 overexpressing neurons (Fig. 6F). This finding reinforces the link between miR-7 and glutamatergic signaling, indicating that the diminished glutamatergic response attributed to miR-7 is, to some extent, a result of suppression of AMPA signaling achieved through the downregulation of Gria1.

## DISCUSSION

Here we identify oxygen and glucose deprivation as a metabolic stressor triggering endogenous changes in the ncRNA network of ciRS-7 – miR-7 – Cyrano both *in vitro* and *in vivo* during permanent ischemic stroke. In OGD *in vitro* model, we demonstrate dynamic changes in gene expression for these molecules independent of ciRS-7. In *in vivo* condition of transient ischemic stroke, we found no changes in the lncRNA Cyrano. However, the absence of ciRS-7 reduced cellular death and sensorimotor deficits, while the lack of miR-7 resulted in more extensive tissue damage. This suggests that both ciRS-7 and miR-7 may play a more crucial role in this system. In an *in vitro* model recapitulating the molecular changes occurring during ischemic stroke (low ciRS-7 and high miR-7), these effects were partly executed through miR-7-mediated regulation of glutamatergic signaling, contingent on the absence of ciRS-7. These data suggest that ciRS-7 may regulate miR-7 targeting, highlighting an endogenous regulatory role for the ciRS-7/miR-7 network in mediating cellular stress responses under pathophysiological conditions.

In mice, ciRS-7 harbors 130 binding sites for miR-7^9^ and, because of its remarkably high expression in neurons^5,9^, the limited molecules of miR-7 in physiological conditions^5^ are likely associated to ciRS-7, as indicated by our pyramidal excitatory neurons CLIP-seq data and by others^9,10^. Moreover, Cyrano, highly abundant in neurons, harbors a single nearly complementary site for miR-7 which mediates miR-7 degradation^8^. Our data show that OGD induces changes in the neuronal physiological landscape by concomitant decrease in ciRS-7 and Cyrano and subsequent upregulation of miR-7. These changes seem to be independently regulated, as the absence of ciRS-7 did not influence OGD-induced alterations in the gene expression in miR-7 and Cyrano. While the anticorrelation between Cyrano and miR-7 follow established dynamics, with Cyrano triggering miR-7 degradation through TDMD^8^, our study report the induction of miR-7 in the absence of ciRS-7, suggesting that during ischemic stroke the expression of miR-7 does not depend on ciRS-7. Consistently, miR-7 overexpression induces ciRS-7 downregulation and this effect is rescued by overexpressing the Cyrano site that mediates miR-7 degradation through TDMD. Moreover, in instances where miR-7 suppresses ciRS-7 levels, miR-7 upregulation during OGD becomes even more pronounced.

The role of miR-7 in ischemic stroke has been studied *in vivo* with controversial results^44,45^ and our extensive animal cohort revealed substantial biological variability in the expression levels of miR-7 in a mouse model of tMCAo. This outcome is potentially tied to established feedback loops among ciRS-7, Cyrano, and miR-7, as tMCAo in ciRS-7 knockout mice results in a significant overexpression of miR-7 one day post-ischemia, accompanied by a reduced stroke lesion size and motor deficits. Additionally, at 7 dpi, inducible miR-7 knockout mice exhibited an exacerbated ischemic lesion size compared to WT animals, which aligns with literature reporting that miR-7 is necessary for stroke recovery^46^, further suggesting a role for miR-7 in regulating ischemic damage. The transient ciRS-7 KO phenotype is explained by an initial advantage provided by higher miR-7 levels, while subsequent ciRS-7 downregulation in WT animals minimizes genotypical differences at later timepoints. Furthermore, our in vitro data, mimicking molecular changes in ischemic conditions (low ciRS-7, high miR-7), indicate that under physiological conditions highly abundant ciRS-7 may function as a buffering system to regulate miR-7 targeting.

Given the lack of extensive changes in miR-7 targets in both ciRS-7, Cyrano and miR-7 KO animals in physiological conditions^8,9^, the exact role of ciRS-7 as an influencer of miR-7 targeting has remained controversial. Our data show a significant shift in the overall expression of miR-7 targets during OGD, where a concerted downregulation of two known miR-7 regulators, ciRS-7 and Cyrano, occurs. Our data suggest that the inducible nature for this network may explain the reported absence of drastic effects on miR-7 targets in physiological conditions. Consistent with this concept, numerous studies show that single miRNA mutants exhibit a phenotype solely under stress conditions^47,48^. Moreover, unlike experimental settings involving acute treatment, knock-out models are subjected to the buffering effect of cellular adaptation, as previously experienced^9^.

The analysis of Ago2 CLIP-seq neuronal dataset revealed the lead miR-7 targets downregulated in ciRS-7 KO ischemic mice belong to the functional class of glutamatergic signaling and neuronal outgrowth. Glutamate plays a well-established role in ischemic stroke pathophysiology^12^, contributing to neuronal loss through exaggerated release and impaired clearance of glutamate in the synaptic cleft^11^. Intriguingly, ciRS-7 KO neurons exhibit a pronounced sensitivity to glutamate in physiological conditions, which represent a disadvantage in ischemic conditions, due to the susceptibility to glutamate-mediated excitotoxicity. However, ciRS-7 KO neurons do not exhibit increase cellular death upon OGD, but rather present a distinct regulation of processes following OGD response, both *in vitro* and *in vivo*. We propose that the ameliorative effect during ischemic stroke observed ciRS-7 KO mice may be linked to the levels of miR-7 and its unregulated accessibility to targets. In line with this, overexpressing miR-7 dampened the neural glutamatergic response and increased neuronal outgrowth only in ciRS-7 KO neurons, with no effect on the wild-type neurons. A mechanism through which miR-7 partially exerts this role involves targeting the subunit 1 of AMPA receptors (GluR1) by interacting with the 3’ UTR of its mRNA, Gria1, thereby dampening the AMPA response. Our data confirms that miR-7 effects on both glutamatergic signaling and neurite outgrowth are dependent on ciRS-7 levels. In line with our results, the effect of miR-7 on neurite outgrowth was previously reported in a neuroblastoma cell line^41^ that expresses a negligible amount of ciRS-7 even upon differentiation, suggesting a more direct effect of miR-7 in the absence of its targeting regulator.

A recent study associates miR-7 with energy homeostasis in hypothalamic neurons^39^ during challenging energetic conditions similar to ischemic stroke. However, in this context, alterations in miR-7 were not accompanied by changes in ciRS-7 and Cyrano^39^. Combined with our data revealing diverse dynamics in the activation of the ciRS-7 network by various ischemic insults, this indicates that the network response depends on the type, strength, and duration of the stress and the specific cell type.

In summary, we propose a regulatory role for ciRS-7-miR-7 in glutamatergic signaling through miR-7 target genes, hence contributing to the control of post-ischemia neuronal damage. Our data support the hypothesis of a role for ciRS-7 in buffering miR-7 effects against unwanted changes, thus behaving as a “safe-guide” system. This study suggests a role of intracellular network of non-coding RNAs in regulating pathophysiological processes in the brain.

## LIMITATIONS OF THE STUDY

The dynamic nature of multicellular systems, unlike in vitro cultures, results in data incongruencies across different timepoints and stressors. The intrinsic ability of regulatory molecules to induce phenotypic changes even at low doses complicates causality identification in processes involving multiple cellular events, such as disease onset. Growing number of studies revealing non-coding RNA alterations in pathophysiological in vitro systems underscores the necessity for in vivo research to understand the biological relevance of these complex regulatory networks. For a comprehensive analysis of ciRS-7 in ischemic stroke, future studies may benefit from in vivo investigations using Cyrano KO animals and direct rescue experiments on ciRS-7, although these remain technically challenging and time-consuming.

## MATERIAL AND METHODS

### PRIMARY AND IMMORTALIZED CELLS

#### Primary cortical neuron culture preparation and OGD treatment

Primary cortical neurons were prepared from C57BL/6J and C57BL/6N-Cdr1asem1Nikr (Cdr1as KO and their WT counterpart) embryonic day 15 embryonal cortices. After dissection and removal of the meninges, cortices were incubated 15 minutes at 37 °C in a solution of Krebs buffer (0.126 M NaCl, 2.5 mM KCl, 25 mM NaHCO3, 1.2 mM NaH2PO4, 1.2 mM MgCl2, 2.5 mM CaCl2, supplemented with 45mM BSA, 0.8% of 3.85% MgSO4 and 1% Pen/Strep, pH 7.4) and 0.025% (w/v) trypsin (Sigma-Aldrich, T 9201). Tissue was then treated with 0.008% w/v DNaseI (Sigma-Aldrich, DN25) and 0.026% w/v trypsin inhibitor (Sigma-Aldrich, T9003) and centrifuged at 300 x g for 3 minutes. Cell pellet was resuspended in 3ml of DNaseI/Trypsin solution and then diluted in 7ml of Krebs. After centrifugation at 300 x g for 3 minutes the pellet containing embryonic neurons was resuspended in cortical neurons growth media: Neurobasal (Gibco 21103049), B27 Supplement (Gibco, 17504044), 0.2 mM L-glutamine (Lonza, BE17-605E), 0.01 mg/ml Penicillin/Streptomycin (Gibco, 15140122). Cells were plated on Poly-D lysine (Sigma-Aldrich, P6407) freshly precoated plates (50 μg/ml in sterile water plates for 1h at 37 °C and washed in sterile water prior use). Different density was used for 6-well plates (1.8 million cells per well), 48-well plates (125.000 cells per well), 13mm plastic coverslips (30.000 cells per coverslip). After 5 days in culture half of the in cortical neurons growth media was changed to fresh. Experiments were performed 7 days after the isolation day. Cells were maintained in the incubator 37 °C, 5% CO2. For OGD treatment experiments, before hypoxia induction the media was changed to Normoxic (cortical neurons growth media with Neurobasal changed to Gibco A2477501 supplemented with D-Glucose 25mM and Sodium Pyruvate 0.2mM) or OGD (cortical neurons growth media with Neurobasal changed to Gibco A2477501). Normoxic cells were then put back in the incubator, OGD cells were incubated in hypoxic chamber 37 °C, 5%CO_2_, 1%O_2_ (SCI-tive N, Ruskinn Technology). After the OGD timepoint cells were harvested for RNA extraction or subjected to MTT colorimetric test.

#### MTT test

Cell viability was measured from 48-well plates treating the cells with (3-(4, 5-dimethylthiazolyl-2)-2, 5-diphenyltetrazolium bromide), MTT reagent (Sigma-Aldrich, TOX1) diluted with culture media at a final concentration of 120 μM. Triton-X 100 1% v/v (Sigma-Aldrich, X100) treated wells were used as a positive control for this assay. Plates were incubated for 3-5h at +37°C. After, medium was discarded, formazan crystals were dissolved with dimethyl sulfoxide (DMSO) (Sigma-Aldrich, D2650) for 30min at RT in the dark. Absorbance was read at 585nm using Wallac 1420 Victor2 microplate reader (Perkin Elmer). Wells without cells were used as background and subtracted from the absorbance data, all the six technical replicates were plotted for each of the three biological replicate and the data was normalized on normoxic WT neurons or WT neurons infected with GFP only.

#### Neurite length measurement

Neurons were seeded at a density of 125.000 cells/well in 48-well plates. After OGD exposure or lentiviral infection, neurons were imaged at day 7 for 48 hours with IncuCyte® S3 Live Cell Analysis System (Essen BioScience Ltd.) in bright field and green channel live cell images (two 10x magnification images per well). Acquired data were analysed with the Incucyte® Neurotrack Analysis Software Module (Sartorius) considering the average value of the two images taken for each of the nine technical replicates for all the three biological replicates.

#### RNA isolation and qRT-PCR

Total RNA was extracted from primary cells and ipsilateral/contralateral animal cortex using TRIzol™ Reagent (Invitrogen) following the manufacturer’s instructions. 1μl of GlycoBlue™ Coprecipitant (Ambion) was added at the isopropanol step in each sample. RNA was quantified using a Nanodrop 2000 spectrophotometer. 1 μg of total RNA was used for reverse-transcription of mRNAs, circRNA, lncRNAs and pri-miRNA species with High-Capacity cDNA Reverse Transcription Kit (Applied Biosystems) following the manufacturer’s protocol. SYBR Green qPCR Master Mix (High ROX) (Bimake) and custom designed oligos were used to quantify mRNAs, circRNAs, and lncRNAs following the manufacturer’s indications. Mature microRNAs were reverse-transcribed with TaqMan™ MicroRNA Reverse Transcription Kit (Invitrogen) using miRNA-specific primers supplied in the TaqMan® probe kit for qPCR following the manufacturer’s protocol. Mature microRNAs and pri-microRNA PCRs were performed using Maxima Probe/ROX qPCR Master Mix (Invitrogen) and TaqMan® specific probes (Thermofisher) following the manufacturer’s protocol. The result was analyzed with the ΔΔCT method and normalized geometric mean of 2 internal normalization controls (Gapdh and Rplp0) for SYBR green qPCRs and U6 expression for TaqMan® miRNA and pri-miRNA. TaqMan® probes and sequence of the SYBR Green oligonucleotide primers is available in the Supplementary Table S9.

#### Lentivirus vectors and virus generation

Lentiviral vectors LV1-eGFP (control) or LV1-eGFP-miR-7 were generated by subcloning inserts from pAAV_hSYN1-eGFP-miR-7 and pAAV_hSYN1-eGFP (provided by Thomas B. Hansen) inside LV1 (immunodeficiency virus 1 (HIV-1)-based LV-PGK-GFP) backbone by GenScript Biotech Corporation. The generated construct contained HIV-1-LV backbone with hSYN1-eGFP-miR-7 insert instead of PGK-GFPN inserted creating a terminal SmaI (CCCGGG) and a C-terminal ApaI (GGGCCC) flanking restriction sites. The same was performed for the control vector with hSYN1-eGFP insert only. In addition, GenScript Biotech Corporation generated lentiviral vectors, namely LV1-mCherry-CR1 and LV1-mCherry-MUT, for the overexpression of the Cyrano CR1 region involved in miR-7 degradation through TDMD or a mutated form of it. The sequences for these inserts were obtained from the prior publication by Kleaveland et al (2018)^8^ and are accessible on Addgene (https://www.addgene.org/128768/, https://www.addgene.org/128748/). These sequences were then cloned into the previously established LV1-hSYN backbone, with the creation of terminal AgeI (ACCGGT) and C-terminal SalI (GTCGAC) sites.

3rd generation lentiviral particles were produced by the BioCenter Kuopio National Virus Vector Laboratory in Kuopio, Finland. The viral titer was assessed through qPCR serial dilution quantification using eGFP or mCherry ReadyMade™ Primers (IDT) (Supplementary Table S9). Work with the virus vectors was carried out under permission from Finnish National Supervisory Authority for Welfare and Health, Valvira. Cells were infected at day 2 post isolation with MOI 0.5 achieving 80% positive infected cells assessed by GFP or mCherry expression in fluorescent microscope (Supplementary Figure 7A,D) at day 7. The amount of overexpression of miR-7 or Cyrano CR1 region was assessed by qPCR as above specified (Supplementary Figure 7B,E).

#### Library preparation OGD cortical neurons

All the samples RNA were isolated with TRIzol™ Reagent (Invitrogen) as specified above. RNA samples were treated with TURBO DNA-free™ Kit (Ambion) following the manufacture’s instruction. RNA integrity was assessed through Agilent Bioanalyzer 2100 system with the Agilent RNA 6000 Nano. The concentration of the samples was established with Qubit™ RNA Extended range kit (Invitrogen). From cortical neurons subjected to OGD we generated 1) library to detect circularRNAs and mRNA and lncRNAs using SMARTer® Stranded Total RNA Sample Prep Kit (Takara Bio USA, Inc.) and 2) library for miRNAs detection utilizing NEBNext® Small RNA Library Prep Set for Illumina (New England Biolabs (UK) Ltd) quality checked and size selected using pippin prep method following manufacturer’s protocol. All the libraries were generated following the manufacturer’s protocol. After generation, the libraries were quantified with Qubit™ High Sensitivity DNA kit (Invitrogen) and by qPCR using KAPA Library Quantification Kit for Illumina® Platforms (Roche). Library size was determined with Agilent Bioanalyzer 2100 system using the Agilent High Sensitivity DNA Kit. Cortical neurons samples to detect circularRNAs, mRNA, and lncRNAs were sequenced paired-end 100 cycles on NovaSeq™ 6000 platform (Illumina) and single-read 75 cycles on NextSeq™ 500 system (Illumina) to detect microRNAs.

#### Calcium imaging

Calcium imaging was performed with murine E15 cortical neurons. The neurons were plated onto PDL-coated circular plastic coverslips (13 mm diameter) in 12-well plate at a density of 30,000 cells/coverslip and kept for 5 days in Neurobasal medium supplemented with B27. To quantify and compare the functional expression of glutamate and GABA receptors in neuronal cultures, we used calcium-imaging technique as previously described^49^. Briefly, neuronal cultures were loaded with the cell-permeable indicator Fluo-4am (Life Technologies, F10471) for 30 min at 37 °C, followed by 10 min washout, and placed in the perfusion chamber mounted on the stage of Olympus IX7010 microscope. Neurons were continuously perfused by basic salt solution (BSS) 3 ml/min containing in mM: 152 NaCl, 10 HEPES, 10 glucose, 2.5 KCl, 2 CaCl2, 1 MgCl2 and pH adjusted to 7.4. Test compounds diluted in the BSS to final concentrations were applied through fast perfusion system (Rapid Solution Changer RSC-200, BioLogic Science Instruments). Cells were imaged with 10x objective using Olympus IX-7010. Excitation wavelength was set as 494 nm, sampling frequency 2 FPS. Glutamate (100 µM with the co-agonist glycine 10 µM promoting activation of NMDA receptors subtype) or GABA (100 µM) were applied for 2 s. Finally, KCl (50 mM) application for 2 s was used to distinguish excitable neurons from possible non-neuronal cells. Fluorescence was detected with the Till Photonics imaging system (FEI GmbH) equipped with a 12-bit CCD Camera (SensiCam) with a light excitation wavelength of 494 nm. Calcium responses to neurotransmitters were evaluated from changes in fluorescence intensity of individual neurons. To this end, regions of interest (ROI) of round shape around the cell body were selected from the whole image with the TILL vision Imaging Software (TILL Photonics GmbH). To distinguish from non-neuronal cells, ROI was taken at the time point corresponding to KCl-induced activation of neurons. The intensity values in each ROI were averaged at each time point to form a fluorescence signal for each neuron and normalized to the baseline level. Signals above 5% of the baseline were included in the analysis.

#### N2a Cell culture

Immortalized murine neuroblastoma cells (N2a) were maintained in DMEM, high glucose, GlutaMAX™ Supplement (Gibco, 31966021) media supplemented with supplemented with 10% Fetal Bovine Serum (Gibco, 10270106) and 1% penicillin and streptomycin in temperature controlled humidified incubator (37 °C, 5%CO_2_). As previously published^50^, for differentiation induction 100.000 cells were plated in a 6-well plate and maintained in complete media supplemented with 20μM retinoic acid (Sigma-Aldrich, R2625) in dimethyl sulfoxide (DMSO) with or without serum starvation (2% FBS instead of 10%) changing half of the media every second day for 5 days before collection.

#### Luciferase assay

Human embryonic kidney 293 cells containing the SV40 T-antigen (HEK293T) were cultured in DMEM, high glucose, GlutaMAX™ Supplement (Gibco, 31966021) medium, supplemented with 10% Fetal Bovine Serum (Gibco, 10270106), and 1% penicillin and streptomycin in a temperature-controlled humidified incubator (37 °C, 5%CO2). For the luciferase assay, 20,000 cells were seeded per well in a 96-well plate. The following day, cells were co-transfected with 100ng of murine Gria1 3’ UTR renilla/luciferase plasmid (MmiT076857-MT06-GC, GeneCopoeia) or miRNA 3’ UTR target control plasmid (CmiT000001-MT06-GC, GeneCopoeia) and 100nM of either miRIDIAN miRNA mimic of mmu-miR-532 (C-310769-01-0002, Dharmacon) as a negative control or mmu-miR-7a-5p (C-310591-07-0002, Dharmacon) using Lipofectamine™ 2000 (11668030, Invitrogen) according to the manufacturer’s protocol. After 48 hours, the luciferase assay was conducted with the Luc-Pair Luciferase Assay Kit 2.0 (LF001-GC, GeneCopoeia), following the manufacturer’s instructions, except for the cell lysis step, which was performed directly on the plate in lysis buffer using three freeze and thaw cycles with dry ice. Luciferase data were normalized to Renilla expression and presented as the Fold Change of Relative Fluorescence between the samples with respect to 3’ UTR vector overexpression alone.

### MOUSE STRAINS AND ANIMAL PROCEDURES

All experiments follow the Helsinki Declaration and guidelines set by the European Commission (European Communities Council Directive, 86/609/EEC) and were approved by the National Animal Experiment Board of Finland. Animals were housed with same sex siblings, in controlled temperature, humidity and light (12 hours light/dark cycle) conditions. Animals had access to ad libitum food. Before the beginning of the animal study mice were divided in single cages. Mice were randomized and all the participants were blinded to the genotype (surgery, MRI acquisition, behavioral test, sample collection and data analysis). Prior to the surgery, we use random number generator (GraphPad Prism quick calcs https://www.graphpad.com/quickcalcs/randomN1/) to randomize the mice into treatment groups. This numbering has been used in crescent order on 3-4 months old male mice that were divided into groups. pMCAO study was performed on 3-4 months old BALB/cOlaHsd male mice (n = 6-8 per timepoint). Cdr1as (ciRS-7) KO tMCAO study was performed on 3-4 months old C57BL/6N-Cdr1asem1Nikr^9^ (Cdr1as KO mice and the WT counterpart), provided by Prof. Dr. Nikolaus Rajewsky, MDC, Berlin, Germany (n = 22-24 per genotype). miR-7 inducible KO tMCAO study was performed on 3-4 months old B6.Cg-miR7a1tm1(fl/fl)ms miR7a2tm1(fl/fl)ms miR7btm1(fl/fl)ms Ndor1Tg(UBC-cre/ERT2)1Ejb/Stf and UBC-cre negative counterpart^39^ provided by Prof. Dr. Markus Stoffel, ETH, Zurich, Switzerland (n = 20-26 per genotype).

#### Genotyping

Before the animal studies, ciRS-7 KO and miR-7 KO mice were genotyped as before^9,39^ except for the tissue dissociation step. Briefly, ear puncture samples were digested in 50 mM Sodium Hydroxide solution for one hour at 95 °C. Once equilibrated to room temperature the samples were neutralized with 1M Tris Hydrochloride pH 8. Two microliters of digested samples were used to run the reaction of PCR with Taq 2X Master Mix (New England Biolabs (UK) Ltd) following manufacturer protocol and using the oligonucleotide primers as specified in Supplementary Table S9. The PCR products were separated on a stained 2.5% or 1.5% Agarose gel for ciRS-7 and miR-7 KO respectively and run in an electrophoresis chamber. Images were acquired with ChemiDoc Imaging Systems (Bio-Rad Laboratories, Inc).

#### Permanent middle cerebral artery surgery (pMCAo)

Permanent middle cerebral artery occlusion (pMCAo) was performed in 3-4 months old BALB/cOlaHsd male mice as previously described^51^. Briefly, mice were anesthetized using 5% isoflurane and anesthesia was maintained with 2% isoflurane. Temperature of the animals was controlled during the surgery with heating blanket and rectal probe (Harvard apparatus). After the skin incision, the left temporal bone was exposed under the temporal muscle, and 1 mm hole was drilled on top of bifurcation in the middle cerebral artery (MCA). The dura was gently removed, and the MCA was lifted and permanently occluded under the bifurcation using a thermocoagulator (Aaron Medical Industries Inc.). Success of the occlusion was confirmed by cutting the MCA above the occlusion site. After the occlusion, the temporal muscle was replaced, and the wound was sutured. The surgery was performed on a total of 20 mice, 3 mice (one per timepoint) died during the surgery and was then not included.

#### miR-7 KO induction with tamoxifen

Before tMCAO surgery, miR-7 inducible KO mice were injected with 2mg of Tamoxifen intraperitoneally once per day for 5 consecutive days at the age of 3 to 4 weeks as before^39^. Recombination was induced in both Cre+ and Cre-littermates. Tamoxifen (Sigma-Aldrich, T5648) was resuspended in 90% corn oil (Sigma-Aldrich, C8267) and 10% pure ethanol and dissolved at 56°C for 30 minutes in the dark. After injection the mice were put on soft food diet and their weight was monitored daily. Mice showing a successful genomic recombination in ear puncture samples two weeks after injection were included in the study (Supplementary Figure S5A). At the end of the study the animals were analyzed for miR-7 abrogation in the brain through RT-qPCR (Supplementary Figure S5B).

#### Transient middle cerebral artery surgery (tMCAO)

Transient middle cerebral artery occlusion (tMCAO) was chosen to continue the in vivo stroke studies as pMCAO in C57BL/6J produces considerably smaller infarcted regions than in the BALB/c background^52,53^. For this reason tMCAO was performed in 3-4 months old male Cdr1as and miR-7 KO and WT mice as previously described^54^. Briefly, induction of anesthesia was performed with 3.5-4% isoflurane in 0.5 L/min of 100% O2-enriched air and maintained at 1-1.5% isoflurane during the surgery. The body temperature maintained at 36 ± 1°C during surgery with a heating pad attached to a rectal probe. A Doppler probe was fixed on the temporal bone over the MCA to monitor blood flow. The right common, external, and internal carotid arteries (CCA, ECA, and ICA) were exposed through an incision in the neck. The arteries were carefully dissected, and a 6-0 silicon coated monofilament of 0.21 mm of diameter (Doccol Corporation) was inserted into the right CCA or left ECA (Cdr1as and mir-7 KO respectively) and led through the ICA until blocking the origin of the MCA. Blood flow blockage was monitored with the Doppler probe, and animals with less than 80% of blood flow decrease were discarded. After one hour of occlusion, the monofilament was withdrawn allowing blood reperfusion. Animals that did not show a correct reperfusion measured by laser doppler were discarded from the study. Incisions were permanently sutured, and the animals were allowed to recover in a temperature-controlled environment for 24 hours. The surgery was performed on a total of 32 WT and 25 Cdr1as KO mice, and 36 WT and 26 miR-7 KO. In the ciRS-7 animal study a total of 7 mice was excluded as: 2 KO and 2 WT mice lacked reperfusion upon surgery, 2 WT and 1 KO mice died during the study. In the miR-7 animal study a total of 12 mice was excluded as: 3 WT and 2 KO lacked reperfusion upon surgery, 4 WT and 3 KO died during the study.

#### Magnetic resonance imaging (MRI)

MRI was performed at 1, 3, and 7dpi using a vertical 9.4 T/89 mm magnet (Oxford instrument PLC) upon anesthesia with 1.8% isoflurane in 30% O2/70% N2O. We acquired twelve slices of 0.8mm thickness per mouse (echo time 40ms, repetition time of 3000ms, matrix size of 128 × 256 and field of view 19.2 × 19.2 mm^2^) and analyzed the first seven images using the Aedes software (http://aedes.uef.fi/) for MatLab program (Math-works). Upon definition of the region of interest (ROI) of contralateral, ipsilateral, and lesion, the lesion volume normalized on oedema was calculated on the first 7 section as: 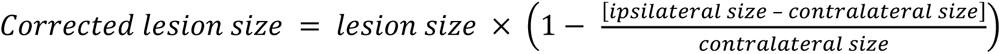. The lesion size is expressed has percentage of lesion on the total brain size.

#### Neurological severity score (NSS)

Mice were examined for neurological deficits at baseline and dpi 1, 3, and 7 using a severity scale comprising the following tests: postural reflex, circling, falling to contralateral side, placement of the contralateral forelimb during motion, and general state of alertness or consciousness. Deficits were graded from 0 (normal) to 2 or 3 (severe)^55^. A sum of these scores were used for statistics. Behavioral assessment was performed by an experimenter blinded to the genotype of mice.

#### Transcardiac perfusion and sample collection

After anesthesia, mice were perfused transcardially with cold saline solution with heparin 2500 IU/l (Leo Pharma A/S). In the pMCAo study, the brains were collected and dissected into contralateral and peri-ischemic cortical regions. In the tMCAO study the brains were cut in 6 coronal sections and stained with 1% 2,3,5-Triphenyltetrazolium Chloride (TTC) (Sigma-Aldrich) in PBS solution for 5 minutes at 37 °C in agitation before dissection of contralateral and peri-ischemic cortical regions. For immunohistochemistry staining, brains were collected and fixed in 4% paraformaldehyde solution in 0.1 M phosphate buffer (PB) pH 7.4. After 22 hours of fixation, the brains were transferred in 30% sucrose in PB buffer solution for 48 hours and then frozen in liquid nitrogen before being stored in - 70 °C.

#### GFAP immunostaining

Each brain was then cut using a cryostat (Leica Microsystems) into six 20 μm coronal sections 400 μm apart, collected on Superfrost™ Plus Microscope Slides (ThermoFisher Scientific) and stored in - 70°C until immunostaining. Sections were then rehydrated with phosphate-buffered saline (PBS) pH 7.4 for 10 min and PBS with 0.05% Tween-20 (PBST) (Sigma-Aldrich) for 5 min. Endogenous peroxidase was blocked by using 0.3 % hydrogen peroxide (H_2_0_2_) in MeOH for 30 min after which sections were washed 3 x 5 min in PBST. Non-specific binding was blocked with 10 % normal goat serum (NGS)(Vector, S-1000) in PBST for 1 h at RT Sections were incubated overnight at RT in primary antibody rabbit anti-GFAP (Agilent, Dako Z0334,1:500 in 5 % NGS-PBST) and then with biotinylated secondary antibody anti-rabbit IgG (H+L) (Vector, BA-1000, 1:200 in 5 % NGS-PBST) for 2 h at RT followed by incubation in ABC reagent (Vector Elite Kit) for 2 h at RT. Sections were washed 3 x 5 min in PBST before and after the incubations. Nickel-3,3’-diaminobenzidine (Ni-DAB) solution (0.175M Sodium acetate, 1% Nickel ammonium sulphate, 50mg DAB (Sigma-Aldrich, D-5905)) with 0.075 % H_2_0_2_ was used to develop the colour for 6 min stopping the reaction by washing the sections 2 x 5 min in dH_2_O. Sections were then dehydrated in 50 % EtOH, 70 % EtOH, 95 % EtOH, 100 % EtOH for 2 min in each and 3 x 5 min in xylene followed by mounting the coverslips with Depex.

#### Image acquisition & analysis

Six GFAP immunoreactivity light microscope images per mouse were acquired by Leica DM6B-Z Thunder Imager microscope (Leica Microsystems CMS GmbH) equipped with DMC2900 camera using 10x magnification. Images were captured using LAS X software (Leica Microsystems CMS GmbH) with exposure time 5 ms, color-gain mode: R:0-G:0-B:25. The images were quantified from the peri-ischemic cortex next to the lesion border (1mm) and the corresponding area of the healthy contralateral hemisphere from 10x images using ImageJ software^56^ function “Measure particles”. The results were presented as relative immunoreactive area to the total area analyzed. This part of the work was carried out with the support of UEF Cell and Tissue Imaging Unit, University of Eastern Finland, Biocenter Kuopio and Biocenter Finland.

#### Library preparation tMCAO tissue samples

All the samples RNA were isolated with TRIzol™ Reagent (Invitrogen) as specified above. RNA samples were treated with TURBO DNA-free™ Kit (Ambion) following the manufacture’s instruction. RNA integrity was assessed through Agilent Bioanalyzer 2100 system with the Agilent RNA 6000 Nano. The concentration of the samples was established with Qubit™ RNA Extended range kit (Invitrogen). We generated a library from ciRS-7 WT and KO tMCAO animals contralateral and peri-ischemic cortices to detect mRNAs changes using CORALL total RNA-Seq Library Prep Kit (Lexogen GmbH) after ribosomal RNA depletion of 600ng of RNA with RiboCop rRNA Depletion Kit V1.2 (Lexogen GmbH). All the libraries were generated following the manufacturer’s protocol. After generation, the libraries were quantified with Qubit™ High Sensitivity DNA kit (Invitrogen) and by qPCR using KAPA Library Quantification Kit for Illumina® Platforms (Roche). Library size was determined with Agilent Bioanalyzer 2100 system using the Agilent High Sensitivity DNA Kit. Animal samples were sequenced single-read 75 cycles on NextSeq™ 550.

### DATA ANALYSIS

#### RNA-seq data analysis

Bulk RNA sequences of mRNA and miRNA have been aligned and quantified to the mouse genome of reference mm10 with the Cdr1as annotation of “chrX:61183248|61186174|.|+” using the and 1.1.0 version of “smrnaseq” applied without clipping and the three prime adapter “AGATCGGAAGAGCACACGTCT”), while circular sequences using 1.1 CIRIquant^57^ set to Read 1 matching the sense strand. Count data were prepared following the workflow defined by Law et al^58^. We filtered out lowly expressed molecules in any condition using the function “filterByExpr” to increase the reliability of the mean-variance relationship. We removed the differences between samples due to the sequencing depth normalizing the count using the trimmed mean of M-values (TMM)^59^ method and applied a log transformation minimizing sum of sample-specific squared difference to enhance the true positive and negative ratio in the downstream analysis^60^. We checked for batch effect due to different timings in biological replicates preparation by performing a principal component analysis and unsupervised consensus clustering with Cola^61^. We identified a batch effect due to different timings in biological replicates preparation influencing the grouping of the normoxic and OGD samples in the dataset ciRS-7 KO/WT cortical neurons (Supplementary Table S5). We corrected the batch effect of this dataset using the negative binomial regression from Combat^62^. We adjusted the variance between the samples as before^63^ through winsorization^64^. We finally created the design matrix for each pair of conditions to compare (contrast) and performed the differential expression analysis using limma/edgeR model^58^ controlling for the false discovery rate with Benjamini-Hochberg Procedure^65^. We employed an interaction term in differential expression analysis to assess whether the OGD response varied between ciRS-7 KO and WT backgrounds. Our analysis using the interaction term identified only one significant DE gene, Gm2004, with a logFC change of 9.631571, aligning with our expectations (Supplementary Table S6). This outcome emphasized that the lack of ciRS-7 did not modulate the OGD response per se and that our backgrounds remained consistent.

#### Ingenuity Pathway Analysis (IPA) and Functional enrichment analysis

We uploaded the differentially expressed genes of each contrast to QIAGEN IPA (QIAGEN Inc., https://digitalinsights.qiagen.com/IPA)^66^ and Metascape^67^ for Ingenuity pathway analysis and functional enrichment analysis, respectively. The analysis has been performed with default parameters and IPA’s background was composed of non-differentially expressed genes.

#### Deconvolution analysis with scRNAseq

Deconvolution aims to estimate the proportions of different cell types within a mixed population in bulk RNA sequencing data using expression profiles from individual cells of scRNA-seq data and represents them in a composite expression profile. The term “contribution” is used to describe the proportion of gene expression attributed to a specific cell type, such as astrocytes or neurons, within the composite profile. We exploited cell-type specific gene expression from external single-cell RNA sequencing (scRNA-seq) data to define the cell subpopulations composing our cortical neuron culture (Supplementary Table S2) and ischemic stroke animal tissue (Supplementary Table S7) bulk RNA sequencing datasets. The scRNA-seq dataset of Loo et al.^68^ of the mouse cerebral cortex at embryonic day 14.5 was provided to SCDC^69^ for performing the deconvolution of our gene expression matrix with cortical neuron samples in normoxic conditions. The scRNA-seq data of Zeisel at al.^70^ composed of murine cerebral cortex samples from p25 to p60 was instead provided to MuSiC^71^ to perform the deconvolution of our gene expression matrix with contralateral and peri-ischemic samples of wild-type and ciRS-7 KO ischemic stroke animals (Supplementary Table S7) as this method is designed to work with multi-subject scRNA-seq dataset.

#### GRO-seq data analysis

A summary of all GRO-seq samples used to quantify pri-miRNA expression levels are presented in Supplementary Table S3. Raw reads for public GRO-seq data were acquired from the GEO database. GRO-Seq reads were trimmed using the HOMER v4.3 (http://homer.salk.edu/homer)^72^ software to remove A-stretches originating from the library preparation. From the resulting sequences, those shorter than 25 bp were discarded. The quality of raw sequencing reads was controlled using the FastQC tool (http://www.bioinformatics.babraham.ac.uk/projects/fastqc)^73^ and bases with poor quality scores were trimmed using the FastX toolkit (http://hannonlab.cshl.edu/fastx_toolkit/). Reads were aligned to mouse mm9 reference genome using the Bowtie^74^ version bowtie-0.12.7. Up to two mismatches and up to three locations were accepted per read and the best alignment was reported. The data was used to create Tag Directories using HOMER. To optimize coverage, a combined tag directory representing all samples of a given cell type (under one or several GSE numbers) was created and used for pri-miRNA quantification using the ‘analyzeRepeats.pl’ command and ‘-rpkm - strand +’ options”.

#### miRWalk analysis

We associated the significantly deregulated miRNAs and differentially expressed genes of wild-type and ciRS-7 KO cortical neurons subjected to OGD (Supplementary Table S2, Supplementary Table S5). We filtered miRNA-Target interactions obtained from 3.0 miRWalk^75,76^ database with 99% probability and located in the 3’ UTR region and included Oip5os1 (Cyrano) as target of mmu-miR-7a-5p^8^. We linked the differentially expressed miRNAs and genes of each contrast by anticorrelation (e.g. miRNA with positive log fold change and significant adjusted probability value is associated to genes with negative log fold change, significant adjusted probability value and targets in miRWalk of the miRNA) focusing on mmu-miR-7a-5p.

#### CDF generation

We tested the assumption of anticorrelation between mmu-miR-7a-5p and its miRWalk gene targets using the empirical Cumulative Distribution Function (eCDF). For each contrast, we compared the eCDF (control function) obtained from the values of log fold change of the non-target genes against the one of the miRNA’s targets and tested their equality with the Kolmogorov–Smirnov test^77^. Genes characterized by longer 3’ UTRs often possess a greater number of potential miRNA binding sites and may display distinct expression characteristics owing to their intrinsically less stable nature. To address this concern, we have incorporated a normalization approach inspired by Kleaveland *et al*. (2018)^8^ in which the relationship between fold change and 3’ UTR length is fitted to a linear function using the 3’ UTR length annotated by Eichhorn *et al.* (2014)^78^. The original fold changes are adjusted by subtracting predicted values, assuming these values exclusively represent expected fold changes due to 3’ UTR length. This subtraction is done as absolute values to adjust the magnitude of the original fold change, based on the assumption that 3’ UTR length, as a gene-specific attribute, is independent of class order. The subtraction aims to eliminate the fold change component attributed to 3’ UTR length and retain the component associated with miRNA targeting. To determine the magnitude and direction of the shift of the eCDF of the targets in respect of the non-targets, we measured the area between the two curves following this formula: ∫ {*F*_*Y*_(*t*) − *F*_*X*_(*t*)} *dt* and the Wasserstein distance^79^. Both the considered targets and non-targets genes passed the filtering by expression, count normalization and participated at the differential expression analysis.

#### CLIP-seq analysis

We collected the HITS-CLIP dataset of Argonaute 2 in pyramidal excitatory neurons produced by Tan et al.^40^ (GSE211552) and replicated the original analysis to map and annotate the genomic regions with a significant read cluster (peak) (Supplementary Table S8). Briefly, we performed a quality control with FastQC (http://www.bioinformatics.babraham.ac.uk/projects/fastqc)^73^ and applied trimming to remove bases after 50 base pairs due to low quality. We applied Cutadapt^80^ to remove adapters and over-represented sequences. We selected reads of at least 24 nucleotides in length and quality score higher than 20 for each nucleotide. Reads were aligned to the mouse reference genome mm10 using Burrows-Wheeler aligner^81^ with default parameters, then the mapped reads were expanded by 30 nucleotides on each side. We considered only reads present in at least 2 experimental replicates and clusters composed of at least 10 overlapping reads. Bowtie^82^ was used to map all occurrences of sequences complementary to the 6mer seed sequence (positions 2-7 of the mature miRNA) of mouse miRNAs from miRBase V21 (https://www.mirbase.org/) to the mouse genome mm10, allowing no mismatches. Strand specific intersection was done with the filtered clusters in the 3’ UTR, 5’ UTR, coding sequence (CDS) and intron regions from RefSeq gene definitions, as well as antisense matches to RefSeq genes^83^. Strand specific intersection was also done to circRNAs from circBase^84^. This generated a genome-wide list with predicted miRNA target sites backed by the detected Ago2 HITS-CLIP clusters for all mouse miRNAs. A subset including all miR-7a-5p targets is supplied as Supplementary Table S8. Only 3’ UTR target sites were used to define miRNA mediated regulation. The full list of all predicted miRNA target sites was uploaded to GEO, as described in the Data Availability section. We conducted quality control on the CLIP-seq output by using MEME-ChIP^85^ (https://meme-suite.org/meme/tools/meme-chip), a dedicated tool for motif enrichment analysis. This tool allowed us to systematically explore the enrichment of miRNA binding motifs within all identified CLIP-seq 3’ UTR clusters. The results of this analysis are provided in Supplementary Table S8.

#### Statistical analysis

Graphs and statistical analysis were performed in GraphPad Prism 9. Every Figure legend reports parameters of replicates (n), statistical test and p-value obtained. Where not specified, p-value was not statistically significant (p-value > 0.05). We refer to n in animal study as single biological replicate (mouse) and in cortical neuron as technical replicates in the same batch, the experiment has been performed in three independent biological replicates.

## Data availability

The data produced and analyzed in this publication have been deposited in NCBI’s Gene Expression Omnibus^86^ and are accessible through GEO Series accession number GSE213179, GSE213067, GSE213177, GSE211552 and GSE215210. The code developed to analyze the data produced in this study is deposited in Zenodo at https://zenodo.org/records/10489728.

## ACKNOWLEDGEMENTS

We thank Nicholas Downes, Mirka Tikkanen, Reetta Vuolteenaho, Miia Salo, Irina Belaia, Nikita Mikhailov, and Gianluca Como for their support. We acknowledge UEF Biocenter Kuopio In Vitro and Ex Vivo Electrophysiology Core Facility, National Virus Vector Laboratory Service, Cell and Tissue Imaging Unit, Lab Animal Centre (Nina Liimatainen and Hanna Miettinen), Biocenter Oulu Transgenic and Tissue Phenotyping Core, Oulu Laboratory Animal Centre Research Infrastructure, Translational Cell Biology Core at Kontinkangas Campus, and the Faculty of Biochemistry and Molecular Medicine of the University of Oulu. This project received co-funding from the Horizon 2020 Framework Programme of the European Union (Marie Skłodowska Curie grant agreement No 740264), as well as the Academy of Finland, Finnish Cultural Foundation, Finnish Foundation for Cardiovascular Research, Instrumentarium Science Foundation, Inkeri and Mauri Vänskä Foundation, Saastamoinen Foundation, Kuopio University Foundation, Aarne and Aili Turusen Foundation, and Aarne Koskelo Foundation. Illustrations were created with BioRender.com.

## AUTHOR CONTRIBUTIONS

F.S. and T.M. conceived and planned the study, with intellectual contributions from T.H., J.J., R. Giniatullin, and D.T. In vivo experiments involved F.S., V.S., P.K., I.U., J.J., H.D., C.P., N.V, and J. Koistinaho. In vitro experiments involved F.S., V.S., D.M.T., M.G.B., R. Giniatullina, N.K., E.G., S.YH. RNA-seq experiments involved F.S., A.H.S., J.S., and J. Kjems, with analysis performed by L.G. Transgenic animals and study assistance were provided by M.L., M.P., M.S., N.R. HITS-CLIP-seq was generated and analyzed by M.V. and A.S. GRO-seq was analyzed by M.K. Sample preparation at Oulu University was done by R.H., F.S., V.S., S.K., and A.H. F.S. and T.M., wrote the manuscript with input from T.H., J.J., R. Giniatullin, J. Kjems, and critical feedback from all authors.

